# DCPS modulates TDP-43 mediated neurodegeneration through P-body regulation

**DOI:** 10.1101/2025.06.13.659508

**Authors:** Yingzhi Ye, Zhe Zhang, Yu Xiao, Chengzhang Zhu, Noelle Wright, Julie Asbury, Yongxin Huang, Weiren Wang, Laura Gomez-Isaza, Juan C. Troncoso, Chuan He, Shuying Sun

**Author notes:** This authors contributed equally: Yingzhi Ye, Zhe Zhang.

## Abstract

The proteinopathy of the RNA-binding protein TDP-43, characterized by nuclear clearance and cytoplasmic inclusion, is a hallmark of multiple neurodegenerative diseases, including amyotrophic lateral sclerosis (ALS), frontotemporal dementia (FTD), and Alzheimer’s disease (AD). Through CRISPR interference (CRISPRi) screening in human neurons, we identified the decapping enzyme scavenger (DCPS) as a novel genetic modifier of TDP-43 loss-of-function (LOF)-mediated neurotoxicity. Our findings reveal that TDP-43 LOF leads to aberrant mRNA degradation, via disrupting the properties and function of processing bodies (P-bodies). TDP-43 interacts with P-body component proteins, potentially influencing their dynamic equilibrium and assembly into ribonucleoprotein (RNP) granules. Reducing DCPS restores P-body integrity and RNA turnover, ultimately improving neuronal survival. Overall, this study highlights a novel role of TDP-43 in RNA processing through P-body regulation and identifies DCPS as a potential therapeutic target for TDP-43 proteinopathy-related neurodegenerative diseases.

## INTRODUCTION

Amyotrophic lateral sclerosis (ALS) and Frontotemporal dementia (FTD) are two closely linked neurodegenerative diseases with a life expectancy of 3-5 years and 3-14 years after symptom onset, respectively^1,2^. TDP-43, a nuclear RNA-binding protein encoded by *TARDBP* gene, is related to ALS/FTD. Many *TARDBP* mutations have been found to be causative in ALS/FTD cases^3,4^. Moreover, TDP-43 pathology without mutation, including nuclear clearance and cytoplasmic inclusion, is found in more than 95% ALS and around 50% FTD cases^5^. Additionally, TDP-43 abnormalities are observed in other neurodegenerative diseases, including Alzheimer’s disease (AD)^6,7^, Parkinson’s disease (PD)^8,9^, and Huntington’s disease (HD)^10^, underscoring its role in a broader spectrum of TDP-43 proteinopathies. Increasing evidence suggests that the loss of nuclear TDP-43 likely precedes cytoplasmic aggregation^11^. Previous studies have shown that TDP-43 LOF in neurons or animal models causes cell death and motor deficits, supporting its contribution to disease progression^12–14^. Therefore, elucidating the mechanisms underlying TDP-43 LOF-mediated neurotoxicity may provide new insights into the pathogenesis of TDP-43-related neurodegenerative diseases and reveal potential therapeutic avenues.

In this study, we performed a survival-based genetic screen using the CRISPR interference (CRISPRi) technology to identify genetic modifiers of TDP-43 LOF-mediated toxicity in human neurons. We found that reduction of the decapping scavenger enzyme (DCPS) mitigates the neurotoxicity induced by loss of TDP-43 through the regulation of processing body (P-body), a cytoplasmic ribonucleoprotein (RNP) granule involved in mRNA decay and translation repression^15^. TDP-43 LOF disrupts P-body homeostasis, leading to increased RNA degradation of transcripts critical for neuronal functions. Similar P-body abnormalities were observed in ALS/FTD patient-derived samples, supporting the disease relevance. Reduction of DCPS rescues P-body integrity and RNA turnover, improving the survival of TDP-43-deficient neurons. These findings identify a novel genetic modifier of TDP-43 mediated neurotoxicity and uncover a new layer of RNP granule dysfunction in TDP-43 related neurodegenerative diseases.

## RESULTS

### CRISPRi screening identifies DCPS as a genetic modifier of TDP-43 LOF-mediated neurotoxicity

To identify genetic modifiers of TDP-43 LOF-mediated toxicity in human neurons, we leveraged the CRISPRi-i^3^Neuron screening system^16^. The engineered human induced pluoripotent stem cell (iPSC) can be differentiated into highly homogenous cortical neurons (i^3^Neuron) upon the dox-inducible Neurogenin-2 (NGN2) expression. Additionally, the iPSC line was also engineered to express dCas9-KRAB, which catalyzes the sgRNA-mediated gene expression repression^16^.

We knocked down TDP-43 in differentiated neurons using short hairpin RNA (shRNA) via lentiviral transduction (Fig. 1a, and Fig. S1a). Reduction of TDP-43 for 10 days significantly decreased neuron survival and caused cell death, measured by propidium iodide (PI), which stains for cells with compromised plasma membrane^17^ (Fig. 1b and Fig. S1b). Cleaved Caspase-3 (CC3), an apoptosis marker^18^, was also significantly increased in TDP-43 knockdown neurons (Fig. 1a). These observations confirm that TDP-43 loss of function induces toxicity in i^3^Neurons.

**Figure 1.**
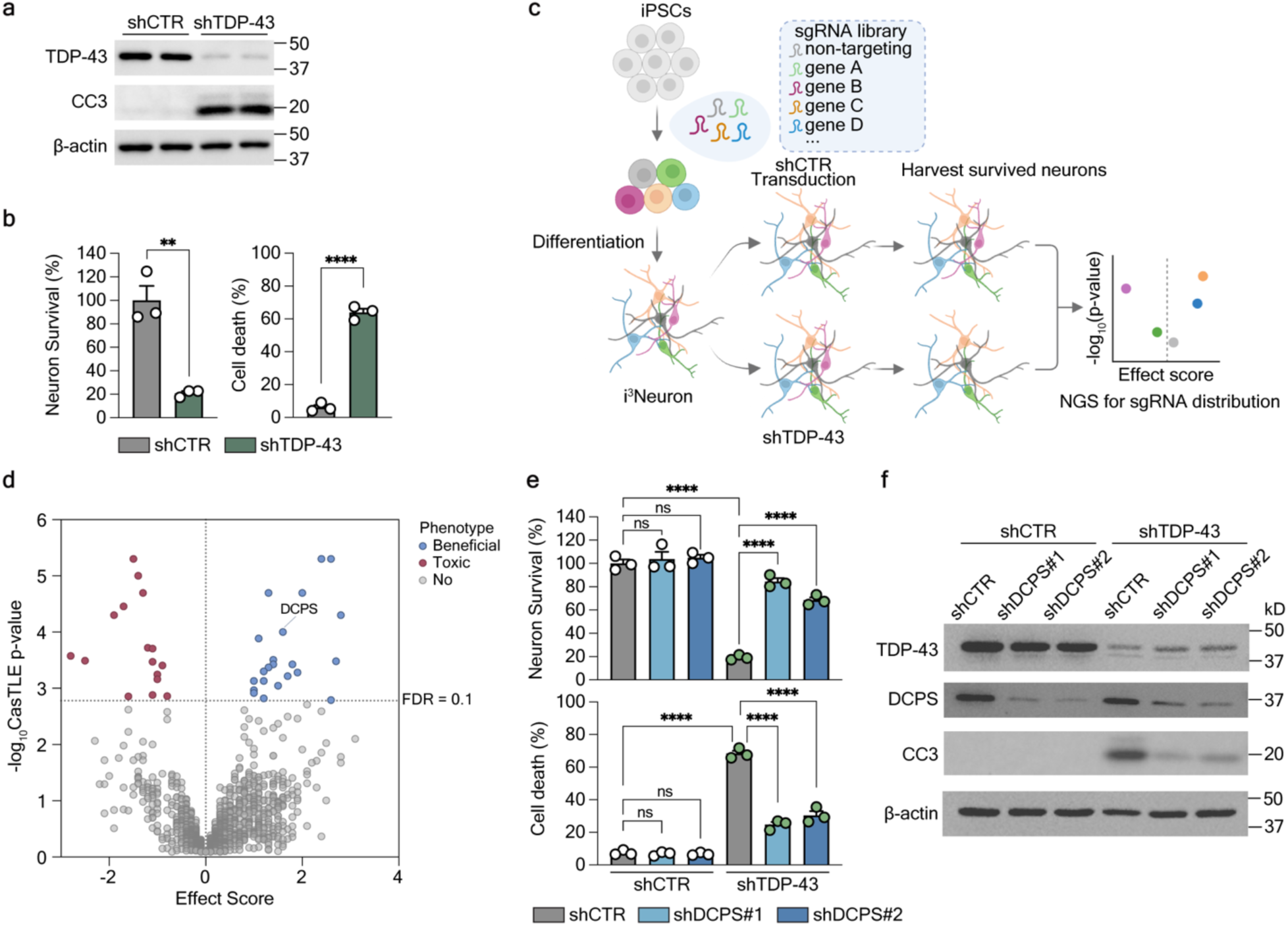
CRISPRi screening identifies DCPS as a genetic modifier of TDP-43 Loss-of-function mediated toxicity in i^3^Neurons. **a**, Western blot shows reduction of TDP-43 and elevation of CC3 in TDP-43 knockdown i^3^Neurons (shTDP-43) on differentiation day 14. β-Actin was used as an internal loading control. n = 2 biological replicates per group. **b**, The percentage of neuron survival and cell death after 10 days of TDP-43 knockdown. Neuron survival was normalized to shCTR group. Cell death was quantified by PI^+^ dead neurons/total neurons. Data are mean ± SEM from three biological replicates. \*\**P* = 0.0033, \*\*\*\**P* < 0.0001, by two-tailed unpaired t test. **c**, The CRISPRi screening scheme. **d**, Volcano plot shows all genes in the CRISPRi screening. Blue: genes promoting the survival of TDP-43 LOF neurons when knocked down. Red: genes enhancing neuron death. Gray: genes with no significant effect. The effect score was calculated from two biological replicates per group by casTLE algorithm. The significant genes were filtered by the cutoff with FDR = 0.1. **e**, Validation of the protective effect of DCPS knockdown (shDCPS #1 and shDCPS #2) in TDP-43 LOF i^3^neurons. Data are mean ± SEM from three biological replicates. \*\*\*\**P* < 0.0001 by one-way ANOVA with Tukey’s post hoc analysis. **f**, Western blot shows CC3 level is decreased when knocking down DCPS in TDP-43 LOF i^3^Neurons. β-Actin was used as an internal loading control.

We then performed the survival-based screening using the CRISPRi druggable library (H1)^19^. The iPSCs were transduced with the pooled lentiviral sgRNA library and then differentiated into neurons. Neurons were infected with lentiviral shRNA targeting TDP-43 (shTDP-43) or non-targeting control (shCTR) on day 4, and the surviving neurons were harvested on day 14^20^ (Fig. 1c). The enrichment of sgRNAs was compared between shCTR and shTDP-43 neurons using the CasTLE algorithm^21^. Genes with enriched sgRNAs in shTDP-43 neurons were considered as beneficial hits, as their downregulation may improve neuronal survival under TDP-43 LOF (Fig. 1d).

DCPS is one of the top beneficial hits, whose reduction can mitigate TDP-43 LOF-mediated toxicity (Fig. 1d). DCPS is a scavenger pyrophosphatase that catalyzes the cleavage of the residual cap structure following 3’ to 5’ RNA degradation. It hydrolyzes the m^7^G-cap from m^7^GpppN to m^7^GMP and NDP^22^. We first validated this hit by designing two individual shRNAs targeting DCPS. We knocked down TDP-43 on day 4 neurons, and followed by DCPS knockdown on day 7 (Fig. S1a). DCPS reduction by both shRNAs (Fig. 1f) significantly mitigated TDP-43 LOF-induced toxicity, as evidenced by improved neuron survival, decreased cell death, and reduced cleaved caspase 3 (Fig. 1e, f and S1c). No obvious toxicity was observed when knocking down DCPS in shCTR neurons (Fig. 1e). We previously found the arginine-rich dipeptide repeat proteins (R-DPRs), poly-GR and poly-PR, produced by *C9ORF72* repeat expansion, induce cell death in i^3^Neurons^20^. DCPS reduction did not decrease R-DPRs-induced toxicity, indicating the protective effect of DCPS knockdown is specific to TDP-43 LOF-mediated neurotoxicity (Fig. S1d).

### DCPS reduction mitigates TDP-43 LOF-induced P-body abnormalities in neurons

To further understand the mechanisms of DCPS-mediated neuroprotection on TDP-43 LOF-induced toxicity, we examined TDP-43 protein levels and found they were unaffected by DCPS knockdown (Fig. 1f), suggesting involvement of other TDP-43 related biological pathways. Given DCPS regulates the m^7^G-cap pool, we used an antibody specific to the m^7^G-cap for immunofluorescence (IF). The m^7^G-cap signals appeared diffuse with small puncta in control neurons, whereas TDP-43 knockdown led to the formation of large cytosolic m^7^G cap granules (Fig. S2a). Notably, DCPS reduction resolved these large m^7^G-cap granules, quantified by decreased m^7^G-cap granule size (Fig. S2a and S2b).

Next, to identify the nature of TDP-43 LOF-induced m^7^G-cap granules, we examined two major cytosolic RNP granules: stress granules and P-bodies. We didn’t observe obvious stress granule formation when knocking down TDP-43, shown by staining of stress granule marker G3BP1 (Fig. S2c). In contrast, m^7^G-cap granules were largely co-localized with the P-body marker DCP1A (Fig. 2a). Consistent with m^7^G-cap granule enlargement, DCP1A granule size was also increased in TDP-43 knockdown i^3^Neurons (Fig. 2a), a finding further confirmed by another P-body marker EDC4 (Fig. 2b and 2c). Moreover, the P-body size was also increased when knocking down TDP-43 in iPSC-differentiated motor neurons (iMNs) (Fig. S2d-f). To further confirm TDP-43 LOF-induced P-body abnormality *in vivo*, we performed EDC4 immunostaining in two TDP-43 conditional knockout (cKO) mouse models, in which TDP-43 was specifically depleted in either ChAT^+^ spinal motor neurons^14^ or CaMKIIα^+^ excitatory neurons in the cortex and hippocampus^23^. P-body size was markedly increased in motor neurons (Fig. 2d, e) and significantly elevated in CaMKIIα^+^ neurons lacking TDP-43 (Fig. S3a, b). There was no stress granule formation (Fig. S3c), suggesting the specificity of P-body changes.

**Figure 2.**
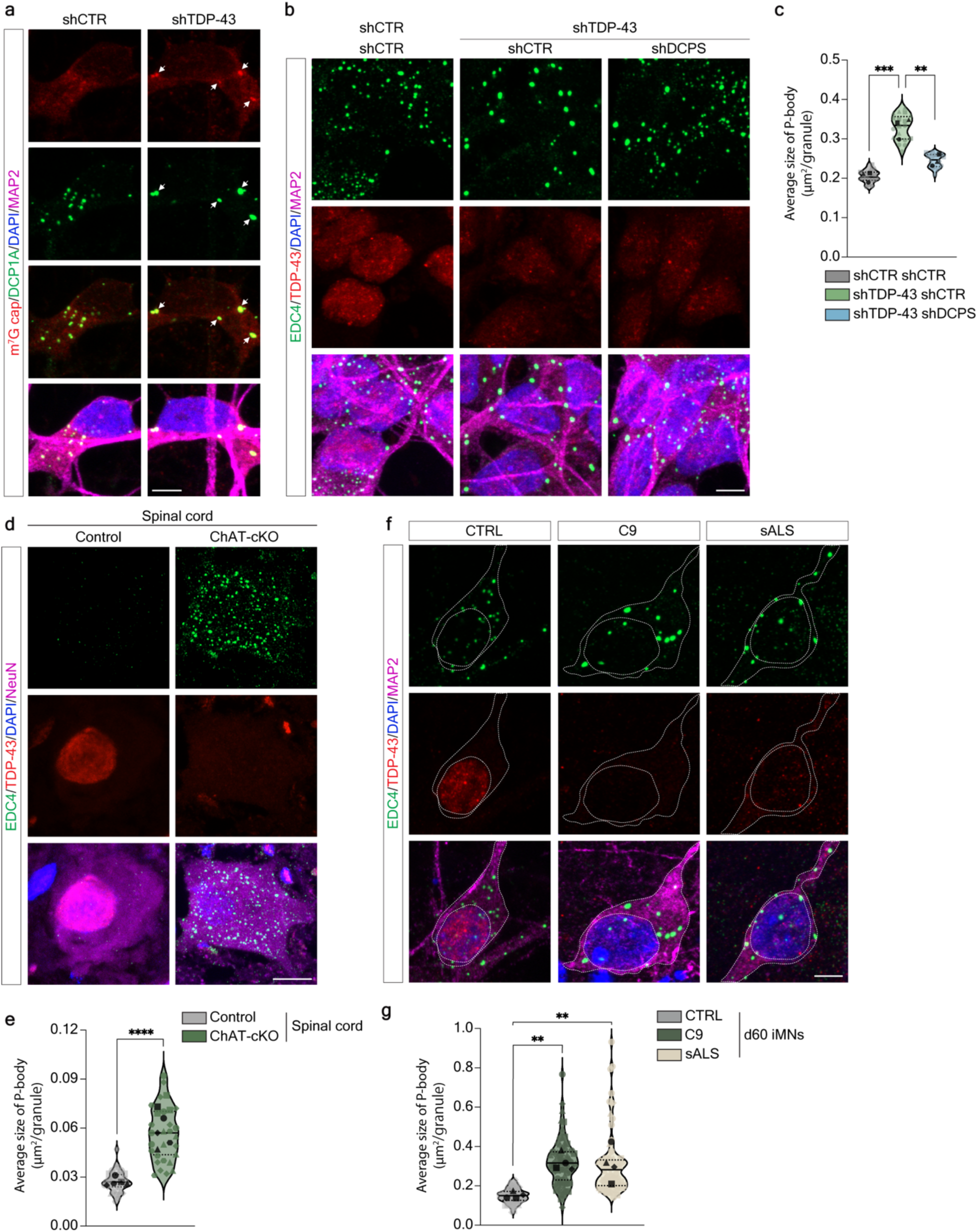
DCPS reduction mitigates TDP-43 LOF-induced P-body abnormality. **a**, Representative images showing co-localization of m^7^G-cap (red) and DCP1A (green) in TDP-43 LOF i^3^Neurons (white arrows). Scale bar: 5 μm. **b**, Representative images of P-body stained by anti-EDC4 (green) in i^3^Neurons. Scale bar: 5 μm. **c**, Quantification of P-body size in (b). The average P-body size was calculated by total granule area/total granule number per imaging field (60×). Each light-colored symbol represents one imaging field. Different shapes represent biological replicates. Black symbols represent the mean value of each biological replicate. Each imaging field included at least 10 cells. 18 imaging fields were quantified per group. Data are median (solid line) with quartiles (dashed line) from three biological replicates. \*\*\**P* = 0.0005, \*\**P* = 0.0041, by one-way ANOVA with Tukey’s post hoc analysis. **d**, Representative images of P-bodies in motor neurons from ChAT^+^ TDP-43 cKO mice (ChAT-Cre; Tardbp^flx/flx^) compared to control mice (ChAT-Cre; Tardbp^flx/+^) at 3-month-old age. Scale bar: 10 μm. **e**, Quantification of P-body size in (d). Each symbol represents one motor neuron and different shapes represent different mice. The quantified cell number: Control n = 28, ChAT-cKO n = 29. Data are median with quartiles from five mice. \*\*\*\**P* < 0.0001, by two-tailed unpaired t test with Welch’s correction. **f**, Representative images of P-body in d60 iMNs differentiated from iPSCs derived from healthy control (CTRL), *C9ORF72*-ALS/FTD (C9), and sporadic ALS (sALS) patients. Scale bar: 5 μm. **g**, Quantification of P-body size in (f). Each symbol represents one cell and different shapes represent distinct iPSC lines. The quantified cell number: CTRL n = 60, C9 n = 71, sALS n = 78. Data are median with quartiles from four lines. \*\**P* = 0.0051 (left), \*\**P* = 0.0065 (right), by one-way ANOVA with Tukey’s post hoc analysis.

Finally, we examined whether DCPS reduction could reverse the P-body size changes. Intriguingly, DCPS knockdown consistently reduced the size of m^7^G-cap granules and P-bodies that were enlarged by TDP-43 LOF in both i^3^Neurons (Fig. 2b, 2c and S2a, S2b) and iMNs (Fig.S2e and S2f). Collectively, these data demonstrate that DCPS reduction can counteract TDP-43 LOF-induced P-body abnormalities, highlighting a potential mechanism through which DCPS contributes to neuroprotection.

### P-body dysregulation in ALS/FTD patient neurons and postmortem tissue

To assess disease relevance, we examined P-bodies in patient-derived neurons. We cultured C9-ALS/FTD and sporadic ALS (sALS) patient-derived iPSC-differentiated iMNs to day 60, when TDP-43 dysfunction was reported to occur^24^. Patient iMNs showed elevated cryptic exons (Fig. S3d). Cells with TDP-43 mislocalization also exhibited enlarged P-bodies (Fig. 2f and 2g). Moreover, we observed enlarged P-bodies in the temporal cortex of both C9-ALS/FTD and FTD-TDP patients (Fig. S3e and S3f). Taken together, increased P-body size was evident in both patient-derived iMNs and postmortem brain tissues, reflecting a downstream consequence of TDP-43 loss of function.

### TDP-43 regulates P-body homeostasis via interactions with P-body proteins

To further explore how TDP-43 regulates P-body homeostasis, we identified TDP-43 interacting proteins by proximity labeling followed by Mass spectrometry^25^. Though TDP-43 is predominantly localized in the nucleus, it shuttles between nucleus and cytoplasm^26^. TDP-43 has been found to play roles in mRNA trafficking and localized translation in neuronal axons and dendrites, indicating the importance of cytosolic TDP-43 in neurons under physiological conditions^27^. To profile the cytosolic TDP-43 interactome, we expressed APEX2-tagged wild-type TDP-43 (TDP-43^WT^) or TDP-43 with a nuclear localization signal (NLS)-deficient mutant (TDP-43^dNLS^) in i^3^Neurons by lentivirus transduction (Fig. S4a and S4b). Biotin-phenol and H_2_O_2_ were added for biotinylating proteins in proximity to TDP-43. Biotinylated proteins were then pulled down using streptavidin beads and analyzed by mass spectrometry. We identified the cytosolic TDP-43 interactome by comparing the protein abundance between TDP-43^dNLS^ and TDP-43^WT^ (Fig. 3a). Intriguingly, a lot of P-body proteins were enriched in TDP-43^dNLS^-expressing neurons, including components of the decapping and deadenylation complexes, which are essential for P-body formation and function^15^ (Fig. 3a and 3b). We validated the top enriched hit, DCP2, a core enzyme of the decapping complex. Co-immunoprecipitation (co-IP) confirmed the interaction between TDP-43 and the P-body component DCP2 (Fig. 3c). However, TDP-43^dNLS^ was diffusely localized in the cytoplasm and did not form granules or colocalize with P-bodies (Fig. 3d, e and S4c, d). We hypothesize that diffused cytosolic TDP-43 interacts with dispersed P-body components, preventing their condensation and thereby influencing P-body formation and dynamics. Supporting this, proximity ligation assay (PLA)^28^ revealed filiform signals only when neurons were incubated with both anti-Flag (TDP-43^dNLS^) and anti-DCP2 antibodies, indicating an interaction between cytosolic TDP-43 and diffused DCP2 (Fig. 3f). We also validated the interaction with EDC3, a scaffold protein in P-body assembly and co-activator of the decapping complex (Fig. 3f). Altogether, these results suggest that cytosolic TDP-43 binds P-body components and may perturb the equilibrium between their diffused and condensate states, thereby modulating P-body assembly and dynamics.

**Figure 3.**
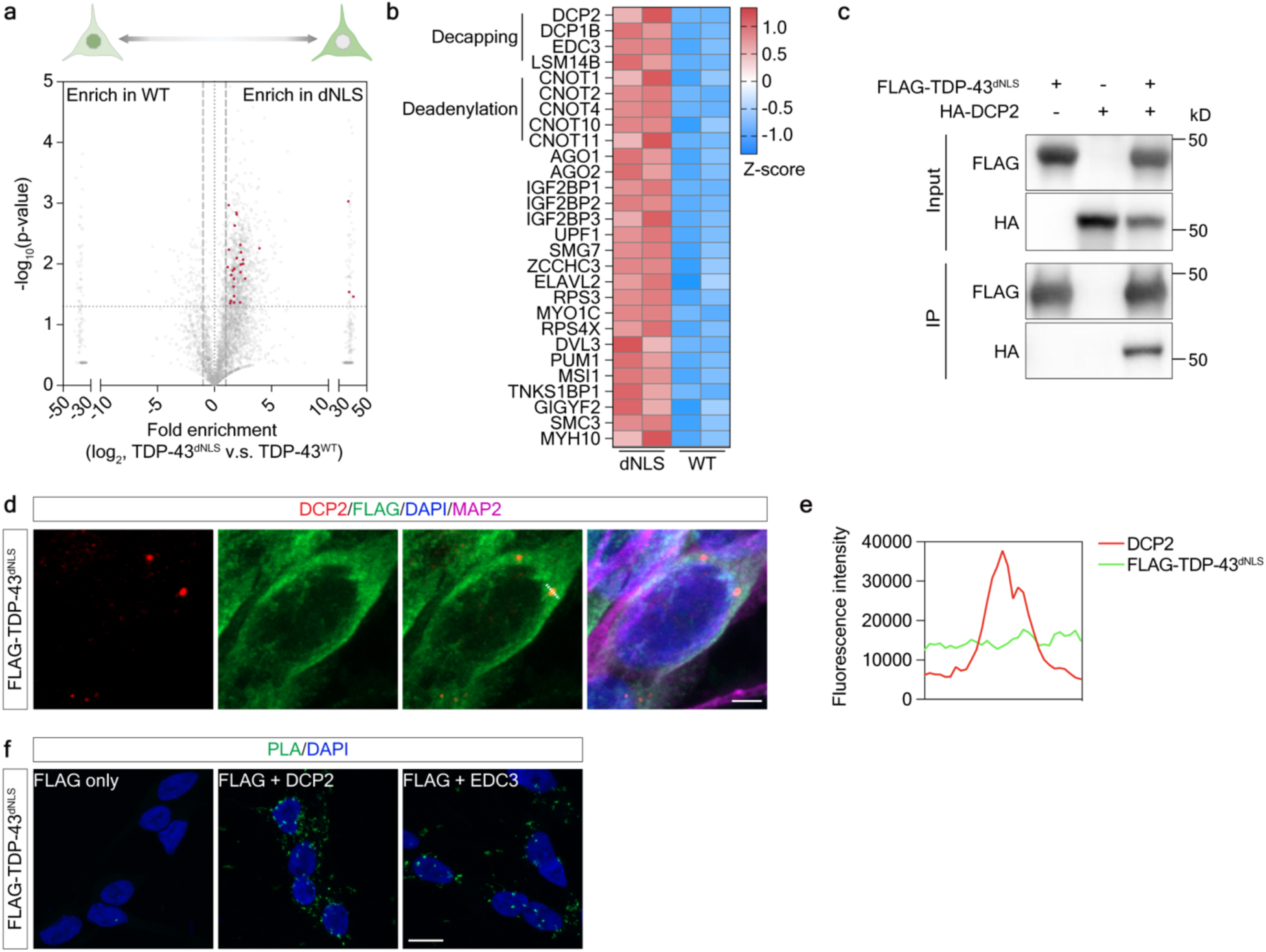
TDP-43 interacts with P-body proteins. **a**, Volcano plot of identified proteins by mass spectrometry shows enriched proteins in TDP-43^dNLS^group (dNLS, right) and in TDP-43^WT^ group (WT, left). Each dot represents a protein. P-body proteins enriched in dNLS are highlighted as red dots (*P* < 0.05 and fold change of protein abundance in dNLS versus WT > 2). *P*-value was calculated by multiple unpaired t-tests from two biological replicates per group. **b**, Heatmap of P-body proteins enriched in dNLS compared to WT. Z-score was calculated by (protein abundance in each sample – mean protein abundance in all samples) / standard deviation. **c**, Co-IP shows TDP-43^dNLS^ and DCP2 interaction in HEK293T cells. IP by anti-FLAG antibody and western blot with anti-HA antibody. **d**, Representative IF images of DCP2 (red) and TDP-43^dNLS^ (green) stained by anti-DCP2 and anti-FLAG antibodies. Scale bar: 2.5 μm. **e**, Line profile of fluorescence intensities in (d). f, Representative images by proximity-labeling assay (PLA). PLA signals (green) show protein-protein interactions. Scale bar: 10 μm.

### TDP-43 LOF-induced P-body alteration dysregulates RNA degradation

We next examined how P-body changes induced by TDP-43 LOF affect the global RNA metabolism. P-body has been indicated to play important roles in RNA degradation and translation^29^. We first examined if TDP-43 knockdown would affect total protein synthesis by puromycin incorporation assay^30^. The newly synthesized proteins can be labeled by puromycin, and then detected by immunoblotting with anti-puromycin antibody. TDP-43 knockdown, with or without DCPS reduction, didn’t show obvious changes in puromycin incorporation (Fig. S5a and S5b), indicating global translation was not affected.

We next measured the transcriptome-wide RNA decay rate to determine potential RNA degradation deficits. The neurons were treated with Actinomycin D to stop de novo transcription, and mRNAs were collected at 0h, 3h and 6h after treatment and subjected to high-throughput sequencing. We observed widespread changes in RNA half-life, particularly downregulation in TDP-43 knockdown neurons (Fig. 4a). Furthermore, downregulated genes strongly correlated with reduced RNA half-life, while upregulated genes showed modest increases in stability (Fig. 4b), supporting that TDP-43 LOF causes profound RNA degradation defects, particularly enhanced decay that contributes to global gene expression downregulation. We compared transcripts with altered stability to previously identified TDP-43 binding targets from individual-nucleotide resolution UV crosslinking and immunoprecipitation followed by RNA-seq (iCLIP-seq)^31,32^. Only a small subset overlapped (Fig. S5c), suggesting that most TDP-43-regulated mRNA degradation occurs indirectly, rather than through direct binding^32^.

**Figure 4.**
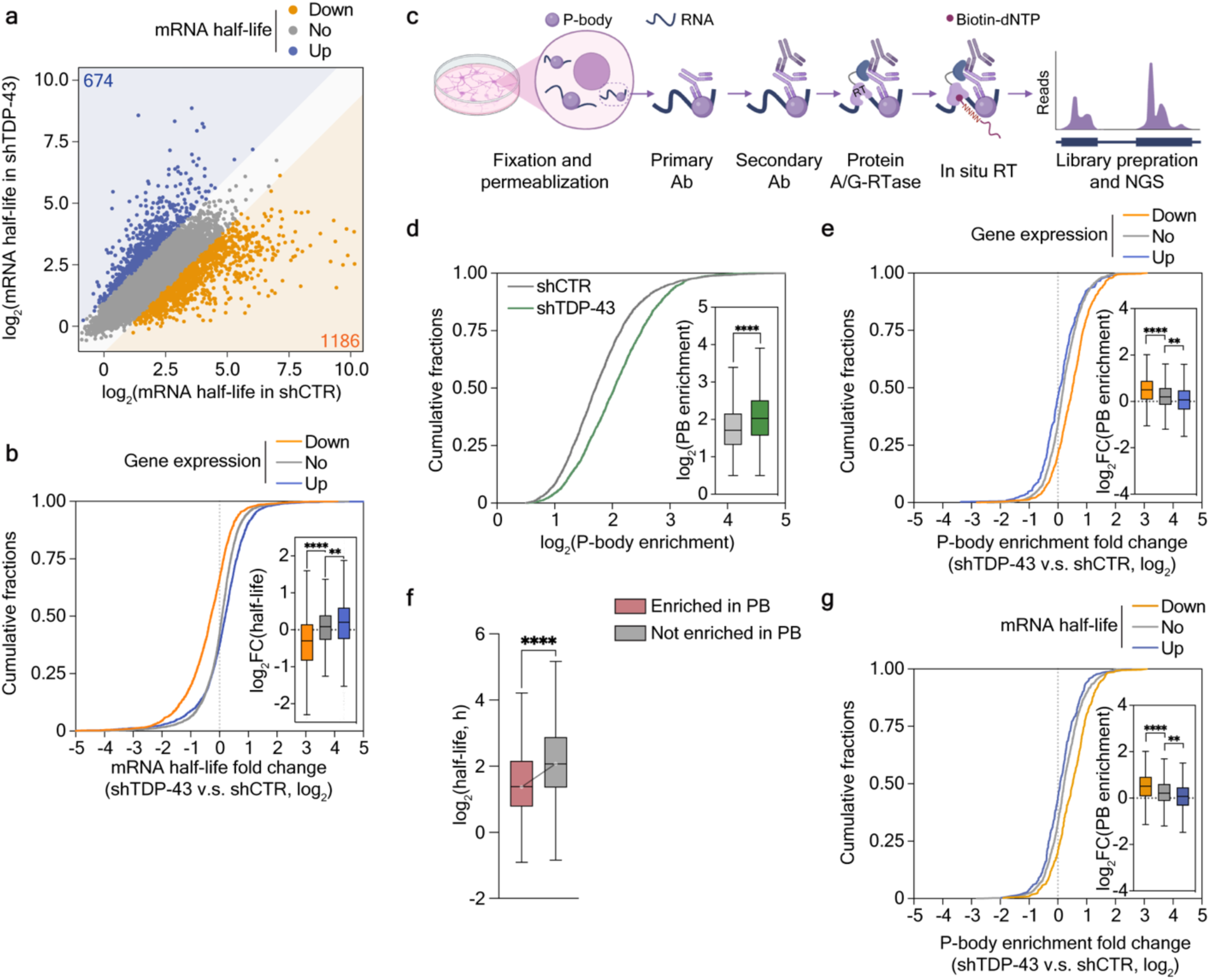
TDP-43 LOF induces P-body and RNA decay deficits. **a**, Scatter plot of mRNA half-life in shTDP-43 (y axis) versus shCTR (x axis) group. Each dot represents a gene. Down (orange): genes whose mRNA half-life is reduced (log_2_FC < −1), n = 1,186. Up (blue): genes whose mRNA half-life is increased (log_2_FC > 1), n = 674. No (gray): genes with no significant changes in mRNA half-life, n=9,821. Genes with mRNA half-life below and equal to zero were filtered out. **b**, Cumulative distribution and box plot (inside) of mRNA half-life changes in shTDP-43 versus shCTR group separated by differential gene expression. Down (orange): down-regulated genes in shTDP-43 compared to shCTR group (log_2_FC < −0.5, adjusted *P* < 0.05), n = 2,423. Up (purple): up-regulated genes in shTDP-43 compared to shCTR group (log_2_FC > 0.5, adjusted *P* < 0.05), n = 2,075. No (gray): genes with no-change in expression level (|log_2_FC| ≤ 0.5 or adjusted *P* ≥ 0.05), n = 7,182. Box plots indicate the interquartile range with the middle line representing the median, and the vertical lines extend to the extreme values. \*\*\*\**P* < 0.0001, by Kolmogorov-Smimov test. **c**, ARTR-seq scheme. **d**, Cumulative distribution and box plot (inside) of P-body enrichment (log_2_ of ARTR-seq peak intensity over input) with or without TDP-43 knockdown. Genes with log_2_ fold > 0.5 and adjusted *P* < 0.05 by DEseq2 (n= 2,857) were included. \*\*\*\**P* < 0.0001, by Wilcoxon matched-pairs signed rank test. **e**, Cumulative distribution and box plot (inside) of ARTR-seq peak intensity changes on P-body enriched genes in shTDP-43 versus shCTR group, stratified by differential gene expression alterations. Down (orange): down-regulated genes in shTDP-43 compared to shCTR group (log_2_FC < −0.5, adjusted *P* < 0.05), n = 647. Up (purple): up-regulated genes in shTDP-43 compared to shCTR group (log_2_FC > 0.5, adjusted *P* < 0.05), n = 298. No (gray): genes with no-change in expression level (|log_2_FC| ≤ 0.5 or adjusted *P* ≥ 0.05), n = 1,912. **** *P* < 0.0001, *** *P* = 0.0004, by Kolmogorov-Smimov test. **f**, Box plot of log_2_ of mRNA half-life of genes enriched (n = 3,971) and not enriched in P-body (n = 7,709) in normal neurons. \*\*\*\**P* < 0.0001, by Kolmogorov-Smimov test. **g**, Cumulative distribution and box plot (inside) of ARTR-seq peak intensity changes on P-body enriched genes in shTDP-43 versus shCTR group, stratified by differential mRNA half-life changes. Down (orange): genes with reduced mRNA half-life in shTDP-43 compared to shCTR group (log_2_FC < −0.5), n = 371. Up (purple): genes with reduced mRNA half-life in shTDP-43 compared to shCTR group (log_2_FC > 0.5), n = 300. No (gray): genes with no-change in mRNA half-life, n = 1,556. Genes with mRNA half-life below and equal to zero were filtered out. **** *P* < 0.0001, ** *P* = 0.0012, by Kolmogorov-Smimov test. **b,d-g**, data are from three biological replicates per group.

To determine whether P-body property changes contribute to mRNA half-life alterations induced by TDP-43 LOF, we examined the P-body associated mRNAs in neurons. We applied the assay of reverse transcription-based RNA-binding protein (RBP) binding site sequencing (ARTR-seq), a recently developed method that can capture dynamic RBP-RNA interactions with high sensitivity and specificity^33^. ARTR-seq couples IF with in situ reverse transcription using biotinylated dNTPs at RBP binding sites, enabling enrichment and sequencing of labeled cDNA that represent RBP-associated RNAs (Fig. 4c). This method has been successfully used to identify RNAs enriched in nuclear speckles^34^, and is well suited for profiling mRNAs associated with P-bodies. We used an antibody targeting the RNA helicase DDX6, a canonical P-body marker and essential RBP for P-body assembly^35,36^. DDX6 immunostaining recapitulated the increased P-body size in TDP-43 LOF neurons, which was rescued by DCPS knockdown (Fig. S5d and S5e). ARTR-seq targeting DDX6 identified 4,483 enriched transcripts in neurons, significantly overlapping with previously published P-body transcriptome by fluorescence activated particle sorting (FAPS) in HEK293 cells^35^, confirming that DDX6 ARTR-seq specifically captured P-body-associated RNAs (Fig. S5f).

TDP-43 knockdown increased overall RNA association with P-bodies (Fig. 4d). Intriguingly, genes whose expression was downregulated by TDP-43 LOF exhibited increased P-body enrichment (Fig. 4e). Consistent with previous findings that rapid RNA decay occurs in P-bodies^37^, P-body-enriched transcripts displayed significantly shorter RNA half-lives compared to non-enriched ones (Fig.4f). Similarly, mRNAs with reduced half-lives in TDP-43 LOF neurons showed increased association with P-bodies (Fig. 4g), supporting that P-body enlargement upon TDP-43 reduction promotes excessive RNA decay by sequestering and accelerating degradation of associated transcripts.

It is generally recognized that 3’UTR length inversely correlates with mRNA stability^38^. In line with this, we found that P-body-enriched genes tend to have longer 3’UTR (Fig. S5g). Likewise, TDP-43 LOF-induced downregulated genes also have longer 3’UTR compared to genes that have no change in expression level (Fig. S5h). These consistent negative correlations between 3’UTR length and RNA half-life in both P-body-associated and TDP-43 LOF-downregulated transcripts further support that TDP-43 and P-body cooperate to fine-tune RNA degradation. Notably, while some TDP-43 LOF-induced cryptic-spliced isoforms are targeted by nonsense-mediated decay (NMD)^39^, they account for only a small proportion of downregulated genes (Fig. S5i), indicating that TDP-43 regulates RNA degradation largely independent of its role in splicing. Altogether, these results show that TDP-43 LOF alters P-body properties, leading to increased mRNA association and accelerated decay, especially of transcripts with long 3’ UTRs. This drives widespread transcriptomic dysregulation, revealing a previously unrecognized role of TDP-43 in modulating RNA stability via P-body dynamics in neurodegeneration.

### DCPS reduction rescues RNA decay defects and transcriptomic dysregulation

Since DCPS reduction mitigates TDP-43 LOF-induced P-body abnormalities, we asked whether it could also rescue the RNA degradation defects. DCPS knockdown significantly reduced the association of RNA with P-bodies and extended the half-lives of transcripts downregulated by TDP-43 LOF (Fig.5a and 5b). Furthermore, it significantly rescued gene expression dysregulation in TDP-43 LOF neurons (Fig. 5c). Gene ontology analysis showed that the rescued downregulated genes were enriched for neuronal function-related pathways (Fig. 5d). For example, mRNA of *SMARCA5*, a chromatin remodeler highly expressed in the brain, showed increased association with P-body, reduced half-life, and decreased expression under TDP-43 LOF, all of which were rescued by DCPS knockdown (Fig. 5e-g).

**Figure 5.**
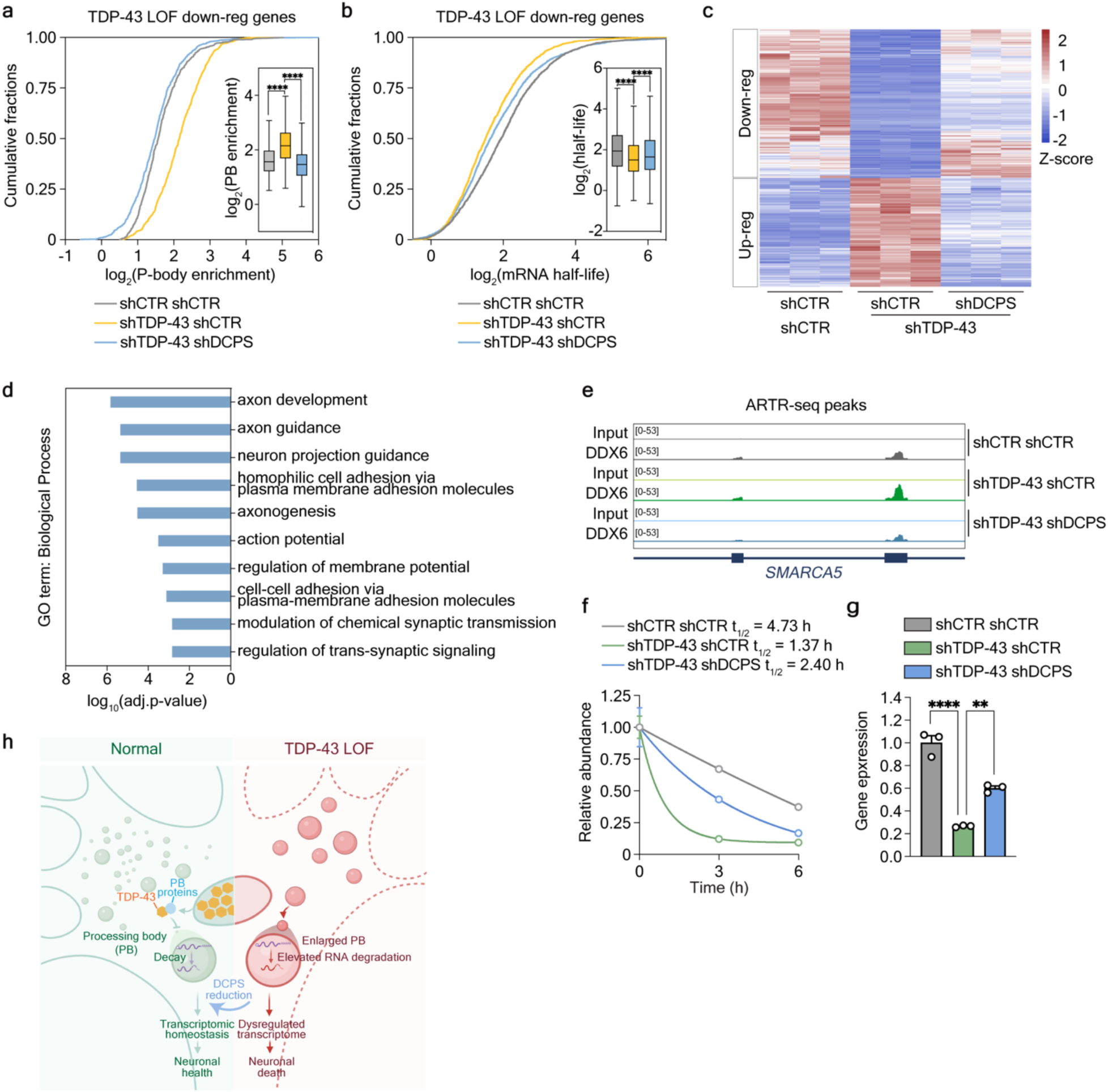
DCPS reduction rescues gene expression dysregulation via correcting P-body and RNA degradation deficits in TDP-43 LOF neurons. **a**, Cumulative distribution and box plot (inside) of ARTR-seq peak intensity changes on P-body-enriched genes that are downregulated in TDP-43 LOF neurons (n = 647) under different conditions. \*\*\*\**P* < 0.0001, by Friedman test with Dunn’s multiple comparisons test. Data are from three biological replicates per group. **b**, Cumulative distribution and box plot (inside) of mRNA half-life changes of genes downregulated by TDP-43 LOF (n = 2423) under different conditions. \*\*\*\**P* < 0.0001, by Friedman test with Dunn’s multiple comparisons test. **c**, Heatmap of gene expression levels of the rescued genes by DCPS reduction in TDP-43 LOF neurons. Downregulated genes induced by TDP-43 LOF that are rescued by DCPS knockdown (log_2_FC < −0.5, adjusted *P* < 0.05 in shTDP-43 shCTR versus shCTR shCTR group and log_2_FC > 0.5, adjusted *P* < 0.05 in shTDP-43 shDCPS versus shTDP-43 shCTR group), n = 929. Up-regulated genes induced by TDP-43 LOF that are rescued by DCPS knockdown (log_2_FC > 0.5, adjusted *P* < 0.05 in shTDP-43 shCTR versus shCTR shCTR group and log_2_FC < −0.5, adjusted *P* < 0.05 in shTDP-43 shDCPS versus shTDP-43 shCTR group), n = 680. Data are from three biological replicates per group. **d**, Gene ontology (GO) term analysis of genes downregulated by TDP-43 LOF and rescued by DCPS knockdown (n = 929) reveals the top 10 biological pathways enriched in neuronal functions. **e**, IGV profiles of DDX6 ARTR-seq peaks on *SMARCA5*. **f**, Half-life of *SMARCA5* mRNA quantified by RNA-seq. Data are mean ± SEM from three biological replicates. **g**, Relative gene expression level of *SMARCA5* quantified by RNA-seq. Data are mean ± SEM from three biological replicates. \*\*\*\**P* < 0.0001, \*\**P* = 0.0018, by one-way ANOVA with Tukey’s post hoc analysis. **h**, A schematic model showing how P-body homeostasis and functions are perturbed by TDP-43 LOF, leading to increased RNA degradation of transcripts critical for neuronal functions. Reduction of DCPS rescues P-body integrity and RNA turnover, improving the survival of TDP-43-deficient neurons.

As TDP-43 has well-known functions in regulating cryptic splicing^40^ and alternative polyadenylation (APA)^41–43^, we also examined whether DCPS knockdown could rescue these RNA processing defects. The analysis showed that DCPS reduction did not reverse TDP-43 LOF-induced changes in cryptic splicing (Fig. S6a-c) or APA profiles (Fig. S6d), reinforcing that DCPS mediates the rescue specifically through the P-body-dependent RNA degradation pathway.

Altogether, these results show that reducing DCPS alleviates TDP-43 LOF-induced defects in P-body RNA composition, RNA degradation, and transcriptomic dysregulation, rescuing neuronal function-related genes and ultimately improving neuron survival (Fig. 5h).

## DISCUSSION

In this study, we employed a survival-based CRISPRi screening and identified DCPS as a potent genetic modifier of TDP-43 LOF-induced neurotoxicity via regulating P-body functions. We found that cytoplasmic TDP-43 interacts with diffused P-body component proteins and modulates P-body formation. This effect may be mediated by influencing the equilibrium between dispersed and phase-separated states of P-body proteins, but the underlying mechanisms require further exploration. Loss of TDP-43 leads to enlarged P-bodies with increased RNA-binding, promoting excessive degradation of associated RNAs and driving widespread transcriptomic dysregulation. These findings uncover a novel function of TDP-43 in regulating RNA stability by modulating P-body dynamics, offering new mechanistic insight into how TDP-43 dysfunction contributes to neurodegenerative disease pathogenesis.

P-bodies are mRNP condensates that function in translation repression and mRNA degradation under physiological conditions^15^, whereas stress granule forms only upon stress stimuli^44^. While stress granules have been extensively studied in neurodegenerative diseases^45^, the importance of P-body homeostasis has remained underappreciated. In Parkinson’s disease (PD), alpha-synuclein aberrantly binds to P-bodies, altering mRNA decay kinetics in patient neurons and brain tissue^46^. Here, we showed that TDP-43 dysfunction also triggers P-body abnormality, highlighting the critical role of P-body dysregulation in the pathogenic mechanisms of TDP-43 proteinopathy-associated neurodegenerative diseases. It is noted that P-body alterations was observed in TDP-43 knockdown HeLa cells^47^, and recent omics studies implicated potential links between TDP-43 and P-bodies^48,49^. This study provides an in-depth mechanistic investigation into P-body dysfunction and its significant contribution to TDP-43 mediated neurotoxicity, highlighting the importance of this previously underappreciated cytoplasmic RNP granule in neurodegeneration.

How P-body size influences its functions was unclear, and mRNAs associated with P-body can be either translationally repressed or targeted for decay. Our findings indicate that increased P-body size in TDP-43 LOF neurons correlates with an overactive degradation of P-body enriched RNAs, rather than influencing global translation efficiency. Additionally, the overall shorter RNA half-lives of P-body-associated transcripts further supports a role for P-bodies in promoting RNA decay in neurons. Notably, recent studies suggest TDP-43 dysfunction compromises the UPF1-mediated RNA surveillance pathway, and particularly altered degradation of transcripts with long 3’ UTRs are implicated in ALS/FTD pathogenesis^50,51^. However, the underlying molecular mechanism remains unresolved. Given that UPF1 has been shown to sense 3’UTR length to guide RNA decay^52^ and our data show that transcripts with longer 3’ UTRs are enriched in P-bodies, it is plausible that P-bodies serve as a platform for facilitating UPF1-mediated decay. Our findings uncover that TDP-43 preserves RNA stability homeostasis by modulating P-body properties and functions in RNA degradation, which likely includes UPF1 substrates along with other RNA surveillance pathways. This provides new mechanistic insights into how TDP-43 dysfunction contributes to transcriptomic disruption in neurodegenerative diseases with TDP-43 proteinopathies.

TDP-43 is known to regulate multiple RNA processing pathways, including cryptic splicing^40^ and APA^41–43^. This study expands the scope of TDP-43’s regulatory functions beyond nuclear RNA processing to cytoplasmic RNA turnover. Importantly, we identified DCPS as a novel genetic modifier of TDP-43 LOF-mediated neurotoxicity. Knockdown of DCPS significantly and specifically improved the survival of TDP-43 deficient neurons, without correcting cryptic splicing and APA alterations, but by restoring P-body function and RNA degradation pathways. Notably, genes with rescued RNA half-lives are enriched in neuronal functions. Altogether, these findings underscore the critical role of RNA surveillance in maintaining neuronal function and health, and highlight an alternative therapeutic revenue of targeting global RNA decay defects, that expands upon current emerging efforts focused on correcting individual cryptic splicing defects. A DCPS inhibitor, RG3039, was previously developed and tested in pre-clinical studies and early clinical trials for spinal muscular atrophy (SMA), a pediatric motor neuron degenerative disease^53–55^. Although the trial was halted due to lack of efficacy, the compound was shown to be well-tolerated and safe. This suggests that reducing DCPS levels is safe and supports the therapeutic potential of repurposing RG3039 for ALS/FTD patients with TDP-43 dysfunction.

This study focuses on the pathogenic mechanisms and disease modifier of TDP-43 LOF-mediated neurotoxicity. While RNA metabolism defects in patients could be more complicated, involving both nuclear clearance and cytosolic aggregates of TDP-43, the immunostaining analyses revealed increased P-body formation in iPSNs and postmortem brain tissues from both *C9ORF72*-linked and sporadic ALS and FTD cases. This provides strong evidence for dysregulated P-body homeostasis in line with loss of TDP-43’s physiological function. Whether pathological or aggregated TDP-43 also influences P-body remains to be determined. As TDP-43 proteinopathy is a common feature across multiple neurodegenerative diseases, these findings may have broader implications for understanding and targeting RNA decay dysregulation in TDP-43 associated pathologies.

## ACKNOWLEDGMENTS

We thank John Hopkins Brain Resource Center for providing postmortem human brain tissue samples. We thank Dr. Philip Wong for sharing the Tardbp^flx/flx^, ChAT-Cre, and CaMKIIα^ERT2^-Cre mice. We thank Dr. Haiyang Yu for sharing the immunostaining protocol for P-body in human postmortem tissues. We thank Dr. Xiaoyang Dou for her guidance in RNA half-life analysis. We thank Dr. Jonathan P. Ling and Irika R. Sinha for providing the list of TDP-43 LOF-induced NMD targets. We thank all the Sun lab members for their helpful discussions. This work is supported by RF1NA127925 and RF1AG078948 from NIH, and Robert Packard Center for ALS Research. Y.Y. received the Toffler Scholar Award. Z.Z. was a recipient of the Milton Safenowitz Post-Doctoral Fellowship from the ALS Association, the Toffler Scholar Award and the Postdoc Development Grant from Muscular Dystrophy Association (MDA). J.A. received Sister Alma Award. C.H. is an investigator of the Howard Hughes Medical Institute.

## AUTHOR CONTRIBUTIONS

Y.Y. and S.S. conceived, designed, and wrote the manuscript with the help of Z.Z. Y.Y. performed most molecular, cell biology, animal experiments, and bioinformatic analysis. Z.Z. helped with DPR experiment, patient line neuron cultures, ARTR-seq experiments, and animal experiments. Y.X. helped with ARTR-seq library construction and sequencing, under the mentorship of C.H. C.Z. helped with patient tissue immunostaining experiments. J.A. and N.W. helped with animal experiments. Y.H. helped with APA analysis and confocal microscopy analysis. W.W. helped with western blot experiments. S.S. supervised the project.

## METHODS

### Plasmids

shRNA constructs were generated following the Addgene protocol (https://www.addgene.org/protocols/plko/). Briefly, the pLKO.1 cloning vector (Addgene #10878) was cut with AgeI and EcoRI and then ligated with annealed oligos containing shTDP-43 or shDCPS. To remove the puromycin resistance gene from the shRNA constructs for use in the puromycin incorporation assay, the PCR fragment containing the blasticidin resistance gene was digested with the NsiI and BamHI and then inserted into the NsiI and BamHI sites of the pLKO.1 vector to replace the original puromycin resistance gene. The following shRNA sequences were used in this study: TDP-43 shRNA (5’-AAGCAAAGCCAAGATGAGCCT-3’), DCPS shRNA#1 (5’-GCTCGATGACTTGTACTTGAT-3’), DCPS shRNA#2 (5’-GCAGTTCTCCAATGATATCTA-3’).

For Lenti-Flag-TDP-43^dNLS^, the PCR fragment containing Flag-TDP-43^dNLS^ sequence (a kind gift from Haiyang Yu) was digested with AgeI and BamHI, and then inserted into the AgeI and BamHI sites of Lenti-GFP-puro vector^21^.

To generate APEX2 constructs used for TDP-43 interactome, PCR fragments containing Flag-TDP-43^WT^ or Flag-TDP-43^dNLS^ and the PCR fragment containing APEX2 were inserted into the AgeI and BamHI sites of Lenti-GFP-puro vector using Gibson assembly (New England Biolabs, E2621S). Flag-TDP-43^WT^ was amplified from pEGFP-N1-TDP-43^56^.

Plasmids for co-immunoprecipitation of TDP-43 and DCP2 were generated as follows. The PCR fragment containing Flag-TDP-43^dNLS^ or HA-DCP2-V5 was digested with AgeI and NheI, and then inserted into the AgeI and NheI sites of the pcDNA3.1 vector. The HA-DCP2-V5 sequence was amplified from pT7-EGFP-C1-HsDCP2 (Addgene #25031).

### Human cell lines

HEK293T cells were cultured in DMEM (Sigma-Aldrich, Cat. No. D5796) supplemented with 10% (v/v) FBS (VWR, Cat. No. 97068-091) and 1× antibiotic-antimycotic solution (Sigma-Aldrich, Cat. No. A5955) at 37°C in a humidified incubator containing 5% CO₂.

### Cell transfection and lentiviral packaging

To package the lentivirus, HEK293T cells were transfected at 60-70% confluency with lentiviral packaging plasmids PRRE (Addgene #12251), PREV (Addgene #12253), pVSVG (Addgene #12259), and co-transfected with lentiviral plasmids using PenFects Transfection Reagent (LifeSct, Cat. No. M0001-01). Virus medium (culture medium plus 1% BSA) was replaced 6 hours post-transfection. Medium containing lentivirus were collected and filtered with 0.45 μm syringe filter (MilliporeSigma, Cat. No. SLHVM33RS) 72 hours post-transfection. The lentivirus was incubated with lenti-X concentrator (Takara, Cat. No. 631232) for at least one hour and then centrifuged at 1,500 g for 45 min at 4℃. Lentiviral pellet was then resuspended in virus medium and stored at –80℃.

Transfection of HEK293T cells to overexpress TDP-43^dNLS^ and DCP2 was performed at 30-40% confluency using TransIT-LT1 (Mirus Bio, Cat. No. MIR2305). pcDNA3.1-FLAG-TDP-43^dNLS^ was co-transfected with pcDNA3.1-HA-DCP2-V5. pcDNA3.1 empty vector was co-transfected with either TDP-43^dNLS^ or DCP2 as a control. Cells were harvested 48 hours post-transfection.

### Human iPSC and iPSC-derived neuron differentiation

Human i^3^N iPSCs and i^3^Neurons differentiation were conducted as previously described^20^. Briefly, engineered iPSCs expressing dCas9 and doxycycline-inducible mNGN2^16^ were maintained in Essential 8 Medium (ThermoFisher Scientific, Cat. No. A1517001) on Matrigel-coated plates (Corning, Cat. No. 354277). Cells were passaged using Accutase (Innovative Cell Technologies, Cat. No. AT104-500) upon reaching ∼80% confluency. To induce neuronal differentiation, iPSCs were dissociated with Accutase and resuspended in N2 medium containing knockout DMEM/F12 (ThermoFisher Scientific, Cat. No. 12660012), N2 Supplement (ThermoFisher Scientific, Cat. No. 17502048), and MEM Non-Essential Amino Acids (ThermoFisher Scientific, Cat. No. 1140050), supplemented with 2 µg/mL doxycycline (Sigma-Aldrich, Cat. No. D9891). After three days of daily medium changes, cells were dissociated and replated on poly-L-ornithine hydrobromide-coated plates (Sigma-Aldrich, Cat. No. P3655) in differentiation medium. The differentiation medium consisted of BrainPhys medium (StemCell Technologies, Cat. No. 05790) supplemented with B27 (Gibco, Cat. No. 17504044), 10 ng/mL NT-3 (ThermoFisher Scientific, Cat. No. 450-03), 10 ng/mL BDNF (ThermoFisher Scientific, Cat. No. 450-02), 1 µg/mL laminin (ThermoFisher Scientific, Cat. No. L2020), and 2 µg/mL doxycycline. Half of the medium was replaced every other day until cells were harvested on day 14.

Peripheral blood mononuclear cell (PBMC)-derived iPSC lines from patients with sALS, C9-ALS, and non-neurological disease controls (Table S1) were obtained from the Cedars-Sinai iPSC Core. iPSCs were maintained and differentiated into spinal motor neurons (iMNs) as previously described (ref). Cells were cultured in mTeSR Plus medium (StemCell Technologies, Cat. No. 100-0276) on growth factor-reduced Matrigel-coated plates (Corning, Cat. No. 354230). Five days post-passaging, differentiation was initiated using S1 medium composed of IMDM (ThermoFisher Scientific, Cat. No. 12440053), F12 (ThermoFisher Scientific, Cat. No. 11765054), NEAA, B27, N2 supplements, penicillin-streptomycin-amphotericin B (PSA; Sigma-Aldrich, Cat. No. A5955), 0.2 µM LDN193189 (Sigma-Aldrich, Cat. No. SML0559), 10 µM SB431542 (StemCell Technologies, Cat. No. 72234), and 3 µM CHIR99021 (Cayman Chemical, Cat. No. 13122). After 6 days, cells were dissociated and re-seeded to S2 medium, which contained all components of S1 with the addition of 0.1 µM all-trans retinoic acid (RA; Sigma-Aldrich, Cat. No. R2625) and 1 µM SAG (Cayman Chemical, Cat. No. 11914). On differentiation day 12, cells were seeded onto Matrigel-coated plates and cultured in S3 medium, consisting of IMDM, F12, NEAA, B27, N2, PSA, RA, SAG, 0.1 µM Compound E (EMD-Millipore, Cat. No. 565790), 2.5 µM DAPT (Sigma-Aldrich, Cat. No. D5942), 0.1 µM db-cAMP (EMD-Millipore, Cat. No. 28745), 200 ng/mL ascorbic acid (Sigma-Aldrich, Cat. No. A4544), 10 ng/mL BDNF (ThermoFisher Scientific, Cat. No. 450-02), and 10 ng/mL GDNF (ThermoFisher Scientific, Cat. No. 450-10).

### Animal

All mouse procedures using mice were approved by the Johns Hopkins University Animal Care and Use Committee (ACUC). Mice were housed in the home cage with free access to food and water in a room with a 12h light/12 hours dark cycle. To knock out *Tardbp* in ChAT^+^ motor neurons, Tardbp^flx/flx^ mice were crossed with ChAT-Cre; Tardbp^flx/+^ mice, producing ChAT-Cre; Tardbp^flx/flx^ mice (ChAT-cKO) and ChAT-Cre; Tardbp^flx/+^ mice (control)^14^. To knock out *Tardbp* in CaMKIIα^+^ neurons, Tardbp^flx/flx^ mice were crossed with CaMKIIα-Cre^ERT2^; Tardbp^flx/flx^ mice, producing CaMKIIα-Cre^ERT2^; Tardbp^flx/flx^ (CaMKIIα-cKO) and Tardbp^flx/+^ mice (control)^23^. Mice were fed with a tamoxifen containing diet (Inotiv, Cat. No. TD.130859) starting at 6 weeks of age for 6 weeks to induce Cre recombinase activity to knock out *Tardbp* in CaMKIIα^+^ neurons. Balanced number of male and female animals were used for studies.

For tissue harvesting and processing, 3-month-old mice were perfused with DEPC-treated PBS. Spinal cords were fixed in 4% PFA, then processed in 10% sucrose overnight followed by 30% sucrose for 2 days. Cryosections (20 μm) were serial sectioned from cryo-blocks into PBS. Floating sections were post-fixed with 4% PFA overnight before being used for IF staining.

### CRISPRi-Cas9 screening

The CRISPRi v2-h1 (H1) library comprises 13,025 elements, including sgRNAs targeting 2,318 genes encoding kinases, phosphatases, and known drug targets (5 sgRNAs per gene), along with 250 non-targeting control sgRNAs^19^. The library was generously provided by Dr. Michael Ward. Screening was performed as previously described, with minor modifications^20^. Briefly, the H1 library was packaged into lentivirus and used to transduce approximately 20 million iPSCs at a multiplicity of infection (MOI) of 0.4. Infected cells were selected with puromycin and subsequently differentiated into neurons. To maintain screening robustness, cell populations were sustained at a minimum of 1,000-fold library coverage after antibiotic selection and 500-fold coverage through the final neuronal stage. On differentiation day 4, neurons were infected with lentivirus encoding either shTDP-43 or a non-toxic control shRNA (shCTR) at an MOI of 1 and harvested on day 14. Genomic DNA from survived neurons was extracted using the NucleoSpin Blood L Midi Kit (MACHEREY-NAGEL, Cat. No. 740954.20). Library preparation was conducted via PCR using Phusion High-Fidelity DNA Polymerase (ThermoFisher Scientific, Cat. No. F-530XL), followed by gel purification (ThermoFisher Scientific, Cat. No. K0692). Sequencing was performed on the Illumina NextSeq platform at a minimum depth of 250 reads per sgRNA. Two biological replicates per condition (shCTR and shTDP-43) were analyzed using the casTLE^21^ (Cas9 High-Throughput Maximum-Likelihood Estimator) algorithm to identify modifiers of TDP-43 LOF-induced toxicity. sgRNA abundance between conditions was compared, and enrichment was calculated as the log-ratio of guide representation. Gene-level effects were computed from the combined scores of the five sgRNAs targeting each gene.

### Treatments in i^3^Neurons and iMNs

i^3^Neurons were infected with lentivirus to knock down TDP-43 on differentiation day 4, day 5 or day 7, depending on the experiment timeline. Lentivirus to knock down DCPS was added on day 7. For inducible R-DPRs, the lentivirus containing PLX304-TRE3G-GR50 or PLX304-TRE3G-PR50^20^ was added to cells on day 6 with Dox to the differentiation medium. To assess neuronal survival and cell death, images were acquired on day 14 using a Nikon TS-2 microscope. For each biological replicate, at least five fields were imaged at 10× object. Three biological replicates were included per experimental group. Propidium iodide (PI; ThermoFisher Scientific, Cat. No. P1304MP) was added to the culture medium at a final concentration per manufacturer’s instruction, 30 minutes prior to cell harvesting. Bright-field and Cy3 fluorescence channel images were then captured. Quantification of surviving neurons and PI-positive (dead) cells was performed in a blinded manner to ensure unbiased analysis.

For puromycin incorporation assay, the neurons were treated with 10 μg/ul puromycin for 30 min at 37℃ with 5% CO_2_. For RNA half-life measurement, 5 μg/ul Actinomycin D was added to neurons for 0, 3, and 6 hours.

iMNs from a non-neurological disease control (Table S1) were infected with lentivirus to knock down TDP-43 on differentiation day 22, followed by transduction with lentivirus containing shDCPS on day 25. Cells were harvested on day 32 for immunofluorescence analysis. For extended culture, control and patient iMNs were maintained until day 60 and then collected for analysis of TDP-43 LOF-induced cryptic exon analysis and immunofluorescence assays.

### RNA extraction and RT-qPCR

RNA was extracted by TRIzol (ThermoFisher Scientific, Cat. No. 15596026), followed by treatment with TURBO DNase I (ThermoFisher Scientific, Cat. No. AM2238). cDNA was generated using high-capacity cDNA reverse transcription kit (ThermoFisher Scientific, Cat. No. 4368813) according to the manufacturer’s instruction. qPCR was performed with technical duplicates for each sample using LiQuant Universal Green qPCR Master Mix (LifeSct, Cat. No. M0016-05) on the CFX96 optical system (Bio-Rad). GAPDH mRNA was used as internal control. All the primers were listed in Table S2.

### APEX labeling and enrichment of biotinylated protein for mass spectrometry

The APEX labeling was performed as previously described^57^. Briefly, one 10 cm dish of i^3^Neurons were transduced with lentivirus to induce APEX2-tagged TDP-43^WT^ or TDP-43^dNLS^ expression at differentiation day 7. On day 14, cells were incubated with 500 μM biotin-phenol (Iris Biotech GmbH, Cat. No. LS-3500) in fresh differentiation medium for 30 minutes at 37 °C with 5% CO₂. Labeling was initiated by adding hydrogen peroxide (Sigma-Aldrich, Cat. No. H1009) to a final concentration of 1 mM with gentle agitation for 1 minute at room temperature. The labeling medium was removed, and cells were washed three times with quenching buffer containing 5 mM Trolox (Sigma-Aldrich, Cat. No. 238813), 10 mM sodium L-ascorbic acid (Sigma-Aldrich, Cat. No. A7506), and 10 mM sodium azide (Sigma-Aldrich, Cat. No. S2002) in Dulbecco’s phosphate-buffered saline (DPBS), followed by two additional washes with DPBS. Cells were then collected in DPBS for biotinylated protein purification.

Cells were then lysed in 400 μL cell lysis buffer (50 mM HEPES, pH 7.4, 150 mM NaCl, 1% NP-40, 0.1% deoxycholate, 0.05% SDS, 0.1 M NaF, 1 mM EGTA, 2 mM PMSF, 1×Proteinase inhibitor cocktail) supplemented with quenching reagents (5 mM Trolox, 10 mM sodium L-ascorbic acid, and 10 mM sodium azide) on ice for 10 min and centrifuged at 3,000g at 4 °C for 10 min. Supernatants were collected and measured by Bradford assay (ThermoFisher Scientific, Cat. No. 23246). The lysate containing 1 mg proteins was incubated with 80 μL Streptavidin Magnetic Beads (Pierce, Cat. No. PI88816) for 3 hours at room temperature with gentle rotations.

The beads were then washed with a serial of buffers: twice with cell lysis buffer, once with 1M KCl, once with 0.1 M Na_2_CO_3_, once with 2M urea in 10mM Tris-HCl (pH 8.0), twice with cell lysis buffer, and once with ddH_2_O. The proteins were eluted in elution buffer (250 mM Tris-HCl pH 6.8, 250 mM SDS, 400 mM DTT) supplemented with 2mM biotin and 20mM DTT by boiling at 90℃ for 10 min. The eluted proteins were snap-frozen and submitted to BGI America Mass Spectrometry Center for protein identification on the Orbitrap Eclipse Tribrid sequencing platform. The APEX-MS samples include two biological replicates per group.

### Protein extraction and Immunoblotting

Cells were rinsed with PBS and dissociated from the plate. Cell pellets were resuspended in RIPA buffer containing 1× protease inhibitor cocktail (Sigma-Aldrich, Cat. No. 11697498001) and kept on ice for 20 min, followed by 20 min centrifuge at 12,000g. The supernatant was collected and protein concentration was measured by BCA assay (Thermo Scientific, Cat. No. 23225). 20μg of protein was mixed with 5× loading dye and boiled at 90°C for 5 min and loaded onto a 10% SDS-PAGE gel. Wet transfer was performed to transfer protein to the nitrocellulose membrane. The membrane was incubated with 5% skim milk (LabScientific, Cat. No. M-0841) for 1 hour, and with the primary antibody in 5% BSA at 4 °C overnight. Primary antibodies are anti-TDP-43 (proteintech, Cat. No. 10782-2-AP, 1:1000), anti-TDP-43 (Abnova, Cat. No. H00023435-M01, 1:1000), anti-DCPS (Santa Cruz, Cat. No. sc-393226, 1:1000), anti-CC3 (CST, Cat. No. 96645, 1:1000), anti-FLAG (proteintech, Cat. No. 20543-1-AP, 1:1000), anti-HA (CST, Cat. No. 3724, 1:1000), anti-puromycin (Millipore, MABE343, 1:1000), anti-GAPDH (CST, Cat. No. 2118, 1:1000), and anti-β-actin (CST, Cat. No. 3700, 1:1000). Blots were washed with 1× TBST on the second day and probed with secondary antibody (Cityva, Cat. No. NA934, 1:10,000; Cityva, Cat. No. NA931, 1:10,000) in 5% skim milk for 1 hour at room temperature. ECL was used to develop signal on the blot. Images were captured by BioRad ChemiDoc imager.

### Co-immunoprecipitation (co-IP)

Cells were lysed in co-IP lysis buffer (50 mM HEPES pH = 7.9, 150 mM NaCl, 10% glycerol, 1% TritonX-100, 1.5 mM MgCl_2_, 1 mM EGTA, 100 mM NaF) supplemented with 1× protease inhibitor cocktail (Sigma-Aldrich, Cat. No. 11697498001), 2 μg/μL PMSF, and 1 mM DTT on ice for 10 min. Cells were sheered 6 times with 26-gauge syringe to help lysis. Lysates were then centrifuged at 3,000g at 4 °C for 10 min. Supernatants were collected and measured by BCA assay (Thermo Scientific, Cat. No. 23225). 0.5 mg of total protein in 500 μL co-IP lysis buffer was pre-incubated with 10 μg Mouse IgG (Millipore Sigma, Cat. No. NI03) and 10 μL protein G beads (ThermoFisher Scientific, Cat. No. 10004D) at 4℃ for 2 hours with gentle rotations. 10 μL of pre-cleared lysates were saved as input (5%). The rest of lysates were incubated with 10 μg anti-FLAG (Sigma, Cat. No. F1804) at 4℃ overnight. The next day, the lysates were incubated with 10 μL protein G beads (ThermoFisher Scientific, Cat. No. 10004D) at 4℃ for 2 hours with gentle rotations. Beads were washed 6 times with co-IP lysis buffer. Beads were boiled at 90°C for 5 min. Eluted proteins were loaded onto a 10% SDS-PAGE gel according to the immunoblotting protocol as described above.

### Proximity ligation assay (PLA)

PLA was performed using the in situ proximity ligation assay kit (Navinci, NT.MR.100) according to the manufacturer’s instructions. Briefly, cells were washed one time with PBS, fixed with 4% Paraformaldehyde (PFA) in PBS for 15 min, and permeabilized with 0.2% TritonX-100 at room temperature. Cells were then incubated in blocking buffer for 1 hour at 37℃ and followed by primary antibody incubation in primary antibody diluent at 4℃ overnight. The antibodies used were anti-FLAG (Sigma, Cat. No. F1804, 1:2000) with anti-DCP2 (Invitrogen, Cat. No. PA5-115102, 1:200) or anti-EDC3 (proteintech, Cat. No. 16486-1-AP, 1:500). On the next day, cells were washed three times with TBST (Tris-buffered saline supplemented with 0.05% Tween-20) and incubated with probe M1 and R2 in probe diluent for 60 min at 37℃. Cells were washed three times with TBST and incubated with enzyme A diluted in buffer A for 1 hour at 37℃, and followed by two times of wash with TBST and subsequent incubation with enzyme B diluted in buffer B for 30 min at 37℃. After two times of wash with TBST, cells were incubated with enzyme C diluted in buffer C (ATTO 488) for 1.5 hours at 37℃. Cells were then washed one time with TBS, stained with DAPI for 5 min at room temperature, and followed by a final wash of 0.1×TBS. Cells were mounted using ProLong™ Gold Antifade Mountant (Invitrogen, P36934).

### Immunofluorescence staining

Cultured cells were fixed with 4% PFA in PBS for 10 min, permeabilized in 0.2% Triton X-100 for 5 min, and blocked in blocking buffer containing 1% bovine serum albumin (BSA, Sigma-Aldrich, Cat. No. A3294) and 2% goat serum (Sigma-Aldrich, Cat. No. G9023) for 1 hour at room temperature. Cells were then incubated with primary antibodies at 4°C overnight. The primary antibodies include anti-TDP-43 (proteintech, Cat. No. 10782-2-AP, 1:500), anti-TDP-43 (Abnova, Cat. No. H00023435-M01, 1:500), anti-m^7^G-cap (MBL, Cat. No. RN016M, 1:100), anti-G3BP1 (BD, Cat. No. 611126, 1:300), anti-FLAG (Sigma, Cat. No. F1804, 1:2000), anti-DCP2 (Novus Biologicals, Cat. No. NBP1-41070, 1:200), anti-EDC4 (abcam, Cat. No. ab72408, 1:500), anti-DDX6 (Millipore, Cat. No. SAB4200837, 1:500), anti-MAP2 (biosensis, Cat. No. C-1382-50, 1:1000), and anti-NeuN (CST, Cat. No. 12943, 1:1000). On the second day, cells were washed with PBS, and incubated with Alexa Fluor 488/546/647 conjugated secondary antibodies (ThermoFisher Scientific, 1:500). Nuclei were counterstained with DAPI.

For patient postmortem tissue staining, paraffin-embedded temporal cortex sections from C9ORF72-ALS/FTD, sporadic FTD/ALS patients and non-neurological age-matched controls were obtained from Johns Hopkins Brain Resource Center with approval from patient consent (see Table S3 for demographic information). The study using patient samples and data was approved by the Johns Hopkins University School of Medicine Office of Human Subjects Research Institutional Review Board. Tissue sections were warmed at 60℃ for 30 min, then rehydrated from sequential 10-min washes: xylene twice, 100% ethanol twice, 95% ethanol twice, 80% ethanol once, 70% ethanol once, 50% ethanol once, 30% ethanol once, PBS once and finally dH2O once. Antigen retrieval was performed with Tris-based commercial buffer (Vector Laboratories, H-3301-250) at 120℃ for 20 min under high pressure. After cooling down naturally to room temperate, the slides were washed three times for 5 min with 1× PBS. The slides were then permeabilized with 0.2% Triton X-100 for 15 min and incubated in blocking buffer for 1 hour. Slides were incubated in anti-EDC4 (abcam, Cat. No. ab72408, 1:500) diluted in the primary antibody buffer (1% BSA, 2% goat serum and 0.002% Triton X-100 diluted in 1× PBS) overnight at 4 °C. The next day, slides were washed three times for 10 min with 1× PBS and incubated with corresponding Alexa Fluor 488/546/647 conjugated secondary antibodies (ThermoFisher Scientific, 1:500) diluted in 1x PBS for 1 h at room temperature. After being washed three times for 10 min with 1× PBS, the slides were incubated with 0.1% Sudan Black (MP Biomedicals, Cat. No. IC15208810) dissolved in 70% ethanol for 30 min at room temperature. The slides were briefly rinsed with 70% ethanol and washed three times for 5 min with PBS. Nuclei were counterstained with DAPI.

For immunofluorescence of mouse spinal cords, floating sections were transferred to PBS, followed by boiling in Tris-based antigen retrieval buffer (Vector Laboratories, H-3301-250) at 95℃ for 20 min. Sections were cooled down to room temperature and washed three times with PBS for 5 min each. Permeabilizations was performed using 0.2% Triton X-100 for 20 min, followed by incubation in blocking buffer for 1 hour. Sections were incubated with primary antibodies at 4℃ overnight. The primary antibodies include anti-EDC4 (abcam, Cat. No. ab72408, 1:500), anti-TDP-43 (proteintech, Cat. No. 10782-2-AP, 1:500), anti-TDP-43 (Novus Biologicals, Cat. No. NBP1-92695SS, 1:500), anti-HuR (Santa Cruz, Cat. No. sc-5261, 1:200), and anti-NeuN (CST, Cat. No. 12943, 1:1000). On the second day, sections were washed with PBS, and incubated with Alexa Fluor 488/546/647 conjugated secondary antibodies (ThermoFisher Scientific, 1:500). Nuclei were counterstained with DAPI.

### Confocal microscopy and image quantification

Confocal images were captured using a Zeiss 980 Airyscan confocal microscopy. Imaging conditions were kept consistent across all replicas. Z-stack images were taken for all of the images. At least six images were taken with more than 60 neurons for each replica. Image J was used for image analysis, with consistent contract settings applied to all images. P-body size was quantified using the following parameters: size 0.1-4μm^2^, circularity 0.4-1.

### ARTR-seq library preparation

ARTR-seq was performed as previously described with minor modifications^33^. Briefly, cells cultured in imaging chambers were fixed in 1.5% PFA at room temperature for 10 min. The cells were quenched with 125 mM glycine at room temperature for 1 min followed by two washes with Dulbecco’s PBS (DPBS) and permeabilization in 0.5% Triton X-100 in DPBS on ice for 10 min. Cells were then washed two times with DBPS and blocked with blocking buffer (0.1% BSA in DPBS) supplemented with 2U/μL RNase inhibitor (Vazyme, R301-03) at room temperature for 30 min. Cells were incubated with anti-DDX6 (Millipore, Cat. No. SAB4200837, 1:500) diluted with blocking buffer at room temperature for 1 hour. For input samples, cells were incubated with blocking buffer without primary antibody. Next, cells were stained with Alexa Fluor 488 (ThermoFisher Scientific, 1:500) diluted in blocking buffer at room temperature for 30 min and incubated with pAG-RTase (10 nM in blocking buffer) for an additional 30 min. Cells were washed two times with DBPS between each staining steps.

RT reactions were prepared as follows: 2 μM adapter-RT primer (5′-AGACGTGTGCTCTTCCGATCTNNNNNNNNNN-3′), 0.05 mM biotin-16-dUTP (Jena Bioscience, Cat. No. NU-803-BIO16-S), 0.05 mM biotin-16-dCTP (Jena Bioscience, Cat. No. NU-809-BIO16-S), 0.05 mM dTTP/dCTP (ThermFisher Scientific), 0.1 mM dATP/dGTP (ThermoFisher Scientific), 1 U/μl RNaseOUT (ThermoFisher Scientific, Cat. No. 10777019) in DPBS supplemented with 3 mM MgCl_2_. Cells were incubated with the RT reaction mixture at 37 °C for 30 min, then stopped by adding 20 mM EDTA and 10 mM EGTA at room temperature for 3 min.

Cells were digested with 1 mg/ml proteinase K (ThermoFisher Scientific, Cat. No. AM2548) at 37 °C for 1 hour and 50 °C for additional 2 hours. The samples were recovered by phenol-chloroform extraction (pH 8.0) and concentrated by ethanol precipitation. RNA was digested with RNase H (NEB, Cat. No. M0297S) and RNase A/T1 (Thermo Fisher Scientific, Cat. No. EN0551), followed by biotinylated cDNA enrichment using pre-blocked Dynabeads MyOne Streptavidin C1 (ThermoFisher Scientific, Cat. No. 65001). The beads were pre-blocked with 1 μg/μl UltraPure BSA (Thermo Fisher Scientific, Cat. No. AM2616), 1 μg/μl UltraPure Salmon Sperm DNA Solution (Thermo Fisher Scientific, Cat. No. 15632011) and 1 μg/μl Yeast transfer RNA (tRNA) (Thermo Fisher Scientific, Cat. No. AM7119) at room temperature for 30 min. Next, the 3′ cDNA adapter (5′Phos-8N-AGATCGGAAGAGGTCGTGT-3′SpC3) was ligated by T4 RNA ligase 1 (NEB, Cat. No. M0437M) at 25 °C for 16 hours, and cDNA was recovered with the elution buffer of 95% formamide and 10 mM EDTA (pH 8.0) by boiling at 95°C for 10 min, followed by ethanol precipitation. The library was PCR amplified with next-generation sequencing primers, followed by gel purification to select fragments between 180 and 400 bp. Sequencing was performed with three biological replicates per group at the University of Chicago Genomics Facility on an Illumina NovaSeq 6000 platform.

### ARTR-seq analysis

The analysis followed the published script (https://github.com/mingming-cgz/ARTR-seq/tree/main). Briefly, the raw sequencing reads were trimmed with Cutadapt^58^ (version 5.0). Trimmed reads were first mapped to the human rRNA using bowtie2^59^ (version 2.3.5.1) and ones mapped to rRNA were discarded in the downstream analysis. non-rRNA reads were then re-mapped to the human reference genome (hg38) using STAR^60^ (version 2.7.9a). Mapped reads were deduplicated using UMI-tools^61^ (version 1.1.6) and converted to BigWig using deeptools^62^ (version 3.5.1) for IGV visualization. Peaks were called strand-specifically using macs3^63^ (version 3.0.2), given the input libraries as background. All peak regions were merged and counted using subread^64^ (version 2.0.1).

Peak reads were then analyzed by DESeq2^65^ (version 1.40.2). Peaks were first annotated using ChIPseeker^66^ (version 1.36.0). Gene-level enrichment was calculated by normalizing the peak intensity to peak length and relative peak intensity over all peaks in each gene. P-body enriched genes were identified using the following criteria: adjusted *P* < 0.1 and log_2_(gene-level enrichment) > 0.5.

### RNA-seq

RNA-seq libraries were constructed using NEBNext Ultra II Directional RNA Library Prep Kit (NEB, Cat. No. E7765L) with NEBNext Poly(A) mRNA Magnetic isolation Module (NEB, Cat. No. E7490L) according to the manufacturer’s instructions. Briefly, mRNA was isolated from 200 ng total RNA input using oligo(dT)25 magnetic beads and fragmentated into 200-500 bp. After double-strand cDNA synthesis, adaptor ligation and size selection, samples were PCR amplified for 11-12 cycle with index primers. The quantity and quality of the final libraries were assessed by Qubit and Bioanalyzer. RNA sequencing was performed using Illumina NovaSeq platform with 150 bp paired-end reads. Three biological replicates were performed for each group. The total number of sequenced raw reads was 40 million for each sample. Sequencing raw reads were trimmed using trim-galore^67^ (version 0.6.7) and aligned to the human reference genome (hg38) using HISAT2^68^ (version 2.2.1). Aligned reads were deduplicated using picard package (version 2.26.6, https://broadinstitute.github.io/picard/). The number of reads mapped to GENCODE (https://www.gencodegenes.org/, version 38) annotated genes were counted using subread^64^ (version 2.0.1). DESeq2^65^ (version 1.40.2) was used for differential gene expression analysis. Genes with raw counts lower than 10 were filtered out before DESeq2 analysis. Differential expressed genes were defined as those with |log_2_FC| > 0.5 and adjusted *P* < 0.05. Gene Ontology (GO) term analysis was performed using clusterProfiler^69^ (version 4.8.3).

For alternative splicing analysis to identify TDP-43 LOF-induced cryptic exons, rMATS^70^ (version 4.1.1) was used. Deduplicated HISAT2-aligned bam files were used as inputs. Exons with zero inclusion reads in control and high inclusion reads in TDP-43 knockdown samples with FDR < 0.05 and delta PSI (shTDP-43 – shCTR) > 0.1 were defined as cryptic exons. Alternative polyadenylation analysis was performed using PAPA^43^. Differential alternative polyadenylation events were identified as |delta PPAU| > 0.2 and adjusted *P* < 0.05.

### RNA half-life analysis

For RNA-seq to measure RNA half-life, External RNA Control Consortium (ERCC) RNA spike-in control (Thermo Fisher Scientific, 4456740) was added to each sample before poly(A) selection and library preparation.

RNA half-life analysis was performed as previously described^71^. Raw sequencing reads were processed the same as described above with an additional mapping to External RNA Control Consortium (ERCC) RNA spike-in control (Thermo Fisher Scientific, Cat. No. 4456740). Gene-level reads counted by subread^64^ and then normalized to counts per million (CPM). CPM was converted to attomole by linear regression fitting of the RNA ERCC spike-in. RNA half-life was estimated using the previously published method^72^. The concentration of RNA at a given time (dC/dt) is proportional to the constant of RNA decay (*K*_decay_) and the RNA concentration (C), leading to the following equation:

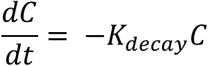

The RNA degradation rate *K*_decay_ was estimated by:

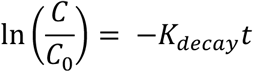

To calculate the RNA half-life (*t*_1/2_), when 50% of the RNA is decayed 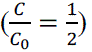, the equation was:

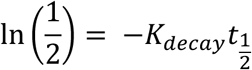

So,

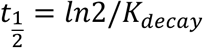

The final RNA half-life was calculated by using the average value of 0, 3, and 6 hours.

### Statistical analysis

Quantification methods of western blots and images have been described in figure legends. The value of n and what n represents is indicated in each figure legend. Analyzed data was plotted and tested for statistical significance using the GraphPad Prism software and RStudio. *P* value of < 0.05 was considered to be significant (\**P* < 0.05, ** *P* < 0.01, *** *P* < 0.001, **** *P* < 0.0001). For each quantification, the type of error bar used and statistical test is specified in the figure legends.

## REFERENCES

1. Rooney, J., et al. A multidisciplinary clinic approach improves survival in ALS: a comparative study of ALS in Ireland and Northern Ireland. *Journal of Neurology*, Neurosurgery & Psychiatry 86, 496–501 (2015).

2. Onyike, C.U. & Diehl-Schmid, J. The epidemiology of frontotemporal dementia. International review of psychiatry 25, 130–137 (2013).

3. Liscic, R., Grinberg, L.T., Zidar, J., Gitcho, M.A. & Cairns, N.J. ALS and FTLD: two faces of TDP-43 proteinopathy. European journal of neurology 15, 772–780 (2008).

4. Gendron, T.F., Rademakers, R. & Petrucelli, L. TARDBP mutation analysis in TDP-43 proteinopathies and deciphering the toxicity of mutant TDP-43. Journal of Alzheimer’s Disease 33, S35–S45 (2013).

5. Jo, M., et al. The role of TDP-43 propagation in neurodegenerative diseases: integrating insights from clinical and experimental studies. Experimental & molecular medicine 52, 1652–1662 (2020).

6. Meneses, A., et al. TDP-43 Pathology in Alzheimer’s Disease. Mol Neurodegener 16, 84 (2021).

7. Amador-Ortiz, C., et al. TDP-43 immunoreactivity in hippocampal sclerosis and Alzheimer’s disease. Ann Neurol 61, 435–445 (2007).

8. Yamashita, R., et al. TDP-43 Proteinopathy Presenting with Typical Symptoms of Parkinson’s Disease. Mov Disord 37, 1561–1563 (2022).

9. Chanson, J.B., et al. TDP43-positive intraneuronal inclusions in a patient with motor neuron disease and Parkinson’s disease. Neurodegener Dis 7, 260–264 (2010).

10. Schwab, C., Arai, T., Hasegawa, M., Yu, S. & McGeer, P.L. Colocalization of transactivation-responsive DNA-binding protein 43 and huntingtin in inclusions of Huntington disease. J Neuropathol Exp Neurol 67, 1159–1165 (2008).

11. Sun, M., et al. Cryptic exon incorporation occurs in Alzheimer’s brain lacking TDP-43 inclusion but exhibiting nuclear clearance of TDP-43. Acta neuropathologica 133, 923–931 (2017).

12. Cascella, R., et al. Quantification of the Relative Contributions of Loss-of-function and Gain-of-function Mechanisms in TAR DNA-binding Protein 43 (TDP-43) Proteinopathies. Journal of Biological Chemistry 291, 19437–19448 (2016).

13. Diaper, D.C., et al. Loss and gain of Drosophila TDP-43 impair synaptic efficacy and motor control leading to age-related neurodegeneration by loss-of-function phenotypes. Human molecular genetics 22, 1539–1557 (2013).

14. Donde, A., et al. Splicing repression is a major function of TDP-43 in motor neurons. Acta neuropathologica 138, 813–826 (2019).

15. Luo, Y., Na, Z. & Slavoff, S.A. P-bodies: composition, properties, and functions. Biochemistry 57, 2424–2431 (2018).

16. Tian, R., et al. CRISPR interference-based platform for multimodal genetic screens in human iPSC-derived neurons. Neuron 104, 239–255. e212 (2019).

17. Riccardi, C. & Nicoletti, I. Analysis of apoptosis by propidium iodide staining and flow cytometry. Nature protocols 1, 1458–1461 (2006).

18. Porter, A.G. & Jänicke, R.U. Emerging roles of caspase-3 in apoptosis. Cell death & differentiation 6, 99–104 (1999).

19. Horlbeck, M.A., et al. Compact and highly active next-generation libraries for CRISPR-mediated gene repression and activation. elife 5, e19760 (2016).

20. Zhang, Z., et al. PTPσ-mediated PI3P regulation modulates neurodegeneration in C9ORF72-ALS/FTD. Neuron (2025).

21. Cheng, W., et al. CRISPR-Cas9 screens identify the RNA helicase DDX3X as a repressor of C9ORF72 (GGGGCC) n repeat-associated non-AUG translation. Neuron 104, 885–898. e888 (2019).

22. Milac, A.L., Bojarska, E. & Del Nogal, A.W. Decapping Scavenger (DcpS) enzyme: Advances in its structure, activity and roles in the cap-dependent mRNA metabolism. Biochimica et Biophysica Acta (BBA)-Gene Regulatory Mechanisms 1839, 452–462 (2014).

23. LaClair, K.D., et al. Depletion of TDP-43 decreases fibril and plaque β-amyloid and exacerbates neurodegeneration in an Alzheimer’s mouse model. Acta neuropathologica 132, 859–873 (2016).

24. Baskerville, V., Rapuri, S., Mehlhop, E. & Coyne, A.N. SUN1 facilitates CHMP7 nuclear influx and injury cascades in sporadic amyotrophic lateral sclerosis. Brain 147, 109–121 (2024).

25. Hung, V., et al. Spatially resolved proteomic mapping in living cells with the engineered peroxidase APEX2. Nature protocols 11, 456–475 (2016).

26. Suk, T.R. & Rousseaux, M.W. The role of TDP-43 mislocalization in amyotrophic lateral sclerosis. Molecular neurodegeneration 15, 45 (2020).

27. Ederle, H. & Dormann, D. TDP-43 and FUS en route from the nucleus to the cytoplasm. FEBS letters 591, 1489–1507 (2017).

28. Söderberg, O., et al. Characterizing proteins and their interactions in cells and tissues using the in situ proximity ligation assay. Methods 45, 227–232 (2008).

29. Parker, R. & Sheth, U. P bodies and the control of mRNA translation and degradation. Molecular cell 25, 635–646 (2007).

30. Schmidt, E.K., Clavarino, G., Ceppi, M. & Pierre, P. SUnSET, a nonradioactive method to monitor protein synthesis. Nature methods 6, 275–277 (2009).

31. Huppertz, I., et al. iCLIP: protein–RNA interactions at nucleotide resolution. Methods 65, 274–287 (2014).

32. Tollervey, J.R., et al. Characterizing the RNA targets and position-dependent splicing regulation by TDP-43. Nature neuroscience 14, 452–458 (2011).

33. Xiao, Y., et al. Profiling of RNA-binding protein binding sites by in situ reverse transcription-based sequencing. Nature Methods 21, 247–258 (2024).

34. Wu, J., et al. Dynamics of RNA localization to nuclear speckles are connected to splicing efficiency. Science Advances 10, eadp7727 (2024).

35. Hubstenberger, A., et al. P-body purification reveals the condensation of repressed mRNA regulons. Molecular cell 68, 144–157. e145 (2017).

36. Ayache, J., et al. P-body assembly requires DDX6 repression complexes rather than decay or Ataxin2/2L complexes. Molecular biology of the cell 26, 2579–2595 (2015).

37. Blake, L.A., Watkins, L., Liu, Y., Inoue, T. & Wu, B. A rapid inducible RNA decay system reveals fast mRNA decay in P-bodies. Nature communications 15, 2720 (2024).

38. Matoulkova, E., Michalova, E., Vojtesek, B. & Hrstka, R. The role of the 3’untranslated region in post-transcriptional regulation of protein expression in mammalian cells. RNA biology 9, 563–576 (2012).

39. Mehta, P.R., Brown, A.-L., Ward, M.E. & Fratta, P. The era of cryptic exons: implications for ALS-FTD. Molecular Neurodegeneration 18, 16 (2023).

40. Ling, J.P., Pletnikova, O., Troncoso, J.C. & Wong, P.C. TDP-43 repression of nonconserved cryptic exons is compromised in ALS-FTD. Science 349, 650–655 (2015).

41. Zeng, Y., et al. TDP-43 nuclear loss in FTD/ALS causes widespread alternative polyadenylation changes. BioRxiv (2024).

42. Rot, G., et al. High-resolution RNA maps suggest common principles of splicing and polyadenylation regulation by TDP-43. Cell reports 19, 1056–1067 (2017).

43. Bryce-Smith, S., et al. TDP-43 loss induces extensive cryptic polyadenylation in ALS/FTD. BioRxiv (2024).

44. Protter, D.S. & Parker, R. Principles and properties of stress granules. Trends in cell biology 26, 668–679 (2016).

45. Wolozin, B. & Ivanov, P. Stress granules and neurodegeneration. Nature Reviews Neuroscience 20, 649–666 (2019).

46. Hallacli, E., et al. The Parkinson’s disease protein alpha-synuclein is a modulator of processing bodies and mRNA stability. Cell 185, 2035–2056. e2033 (2022).

47. Aulas, A., et al. G3BP1 promotes stress-induced RNA granule interactions to preserve polyadenylated mRNA. Journal of Cell Biology 209, 73–84 (2015).

48. Xie, L., et al. Context-dependent Interactors Regulate TDP-43 Dysfunction in ALS/FTLD. bioRxiv, 2025.2004. 2007.646890 (2025).

49. Krispin, S., et al. Organellomics: AI-driven deep organellar phenotyping reveals novel ALS mechanisms in human neurons. bioRxiv, 2024.2001. 2031.572110 (2024).

50. Alessandrini, F., Wright, M., Kurosaki, T., Maquat, L.E. & Kiskinis, E. ALS-associated TDP-43 dysfunction compromises UPF1-dependent mRNA metabolism pathways including alternative polyadenylation and 3’UTR length. bioRxiv, 2024.2001. 2031.578311 (2024).

51. Gomez, N., et al. Counter-regulation of RNA stability by UPF1 and TDP43. bioRxiv, 2024.2001. 2031.578310 (2024).

52. Hogg, J.R. & Goff, S.P. Upf1 senses 3′ UTR length to potentiate mRNA decay. Cell 143, 379–389 (2010).

53. Van Meerbeke, J.P., et al. The DcpS inhibitor RG3039 improves motor function in SMA mice. Human molecular genetics 22, 4074–4083 (2013).

54. Gogliotti, R.G., et al. The DcpS inhibitor RG3039 improves survival, function and motor unit pathologies in two SMA mouse models. Human molecular genetics 22, 4084–4101 (2013).

55. Singh, J., et al. DcpS as a therapeutic target for spinal muscular atrophy. ACS chemical biology 3, 711–722 (2008).

56. Cheng, W., et al. C9ORF72 GGGGCC repeat-associated non-AUG translation is upregulated by stress through eIF2α phosphorylation. Nature communications 9, 51 (2018).

57. Wu, R., et al. Disruption of nuclear speckle integrity dysregulates RNA splicing in C9ORF72-FTD/ALS. Neuron 112, 3434–3451. e3411 (2024).

58. Martin, M. Cutadapt removes adapter sequences from high-throughput sequencing reads. *EMBnet*. journal 17, 10–12 (2011).

59. Langmead, B. & Salzberg, S.L. Fast gapped-read alignment with Bowtie 2. Nature methods 9, 357–359 (2012).

60. Dobin, A., et al. STAR: ultrafast universal RNA-seq aligner. Bioinformatics 29, 15–21 (2013).

61. Smith, T., Heger, A. & Sudbery, I. UMI-tools: modeling sequencing errors in Unique Molecular Identifiers to improve quantification accuracy. Genome research 27, 491–499 (2017).

62. Ramírez, F., Dündar, F., Diehl, S., Grüning, B.A. & Manke, T. deepTools: a flexible platform for exploring deep-sequencing data. Nucleic acids research 42, W187–W191 (2014).

63. Zhang, Y., et al. Model-based analysis of ChIP-Seq (MACS). Genome biology 9, 1–9 (2008).

64. Liao, Y., Smyth, G.K. & Shi, W. featureCounts: an efficient general purpose program for assigning sequence reads to genomic features. Bioinformatics 30, 923–930 (2014).

65. Love, M.I., Huber, W. & Anders, S. Moderated estimation of fold change and dispersion for RNA-seq data with DESeq2. Genome biology 15, 1–21 (2014).

66. Yu, G., Wang, L.-G. & He, Q.-Y. ChIPseeker: an R/Bioconductor package for ChIP peak annotation, comparison and visualization. Bioinformatics 31, 2382–2383 (2015).

67. Krueger, F. Trim Galore!: A wrapper around Cutadapt and FastQC to consistently apply adapter and quality trimming to FastQ files, with extra functionality for RRBS data. Babraham Institute (2015).

68. Kim, D., Paggi, J.M., Park, C., Bennett, C. & Salzberg, S.L. Graph-based genome alignment and genotyping with HISAT2 and HISAT-genotype. Nature biotechnology 37, 907–915 (2019).

69. Yu, G., Wang, L.-G., Han, Y. & He, Q.-Y. clusterProfiler: an R package for comparing biological themes among gene clusters. Omics: a journal of integrative biology 16, 284–287 (2012).

70. Shen, S., et al. rMATS: robust and flexible detection of differential alternative splicing from replicate RNA-Seq data. Proceedings of the national academy of sciences 111, E5593–E5601 (2014).

71. Li, Y., et al. Globally reduced N 6-methyladenosine (m6A) in C9ORF72-ALS/FTD dysregulates RNA metabolism and contributes to neurodegeneration. Nature neuroscience 26, 1328–1338 (2023).

72. Liu, J., et al. N 6-methyladenosine of chromosome-associated regulatory RNA regulates chromatin state and transcription. Science 367, 580–586 (2020).

