## Supplementary figures for "DCPS modulates TDP-43 mediated neurodegeneration through P-body regulation"

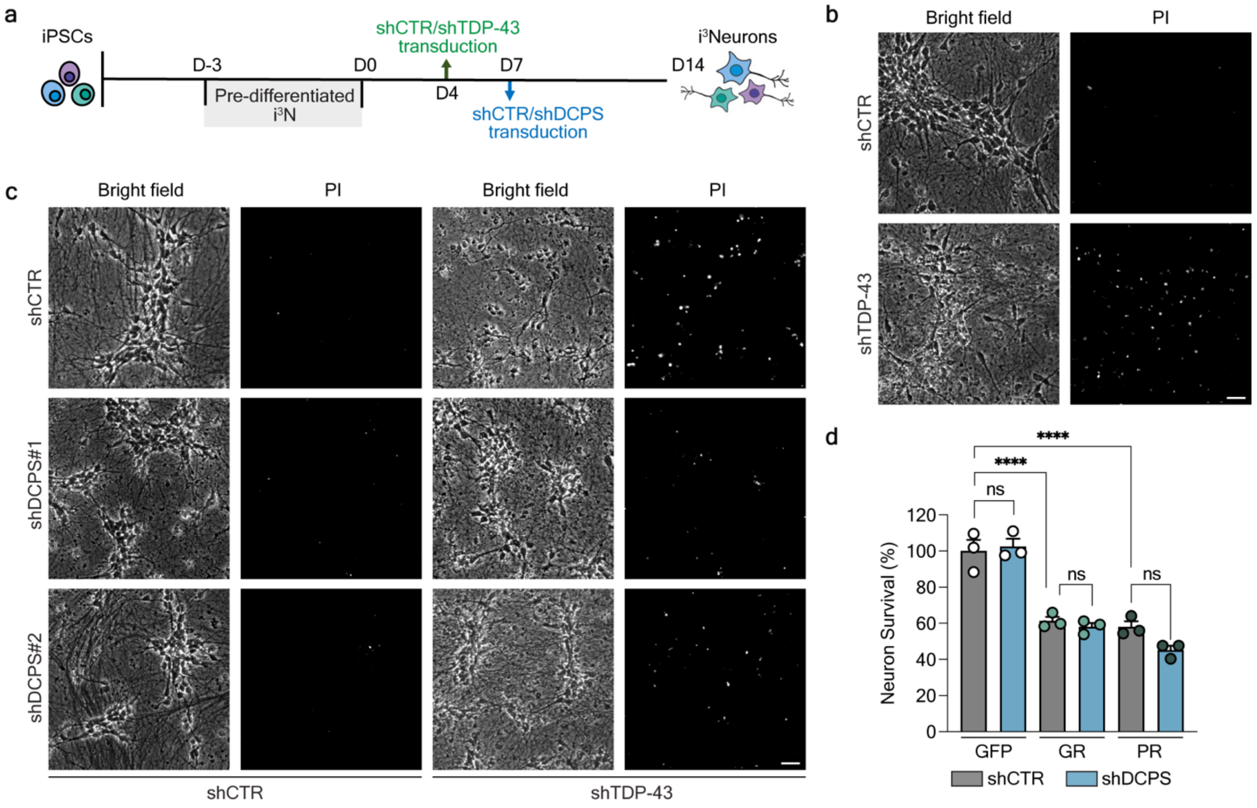


**Figure S1. i$^3$Neuron differentiation and survival assays**

**a**, i$^3$Neuron differentiation diagram and shRNA treatment timeline. **b**, Representative bright field and PI staining images showing significant cell death of TDP-43 LOF i$^3$Neurons (shTDP-43). Scale bar: 20 µm. **c**, Representative images of PI staining showing DCPS reduction rescues TDP-43 LOF-induced neurotoxicity. Scale bar: 20 µm. **d**, Neuron survival quantification to measure R-DPRs-induced toxicity without or with DCPS knockdown. 50-repeats of glycine-arginine (GR) dipeptide, proline-arginine (PR) dipeptide, or GFP control was expressed in i$^3$Neurons by lentiviral transduction on day 6. Neuron survival was normalized to GFP shCTR group. Data are mean ± SEM from three biological replicates. *****P* < 0.0001 by one-way ANOVA with Tukey's post hoc analysis.

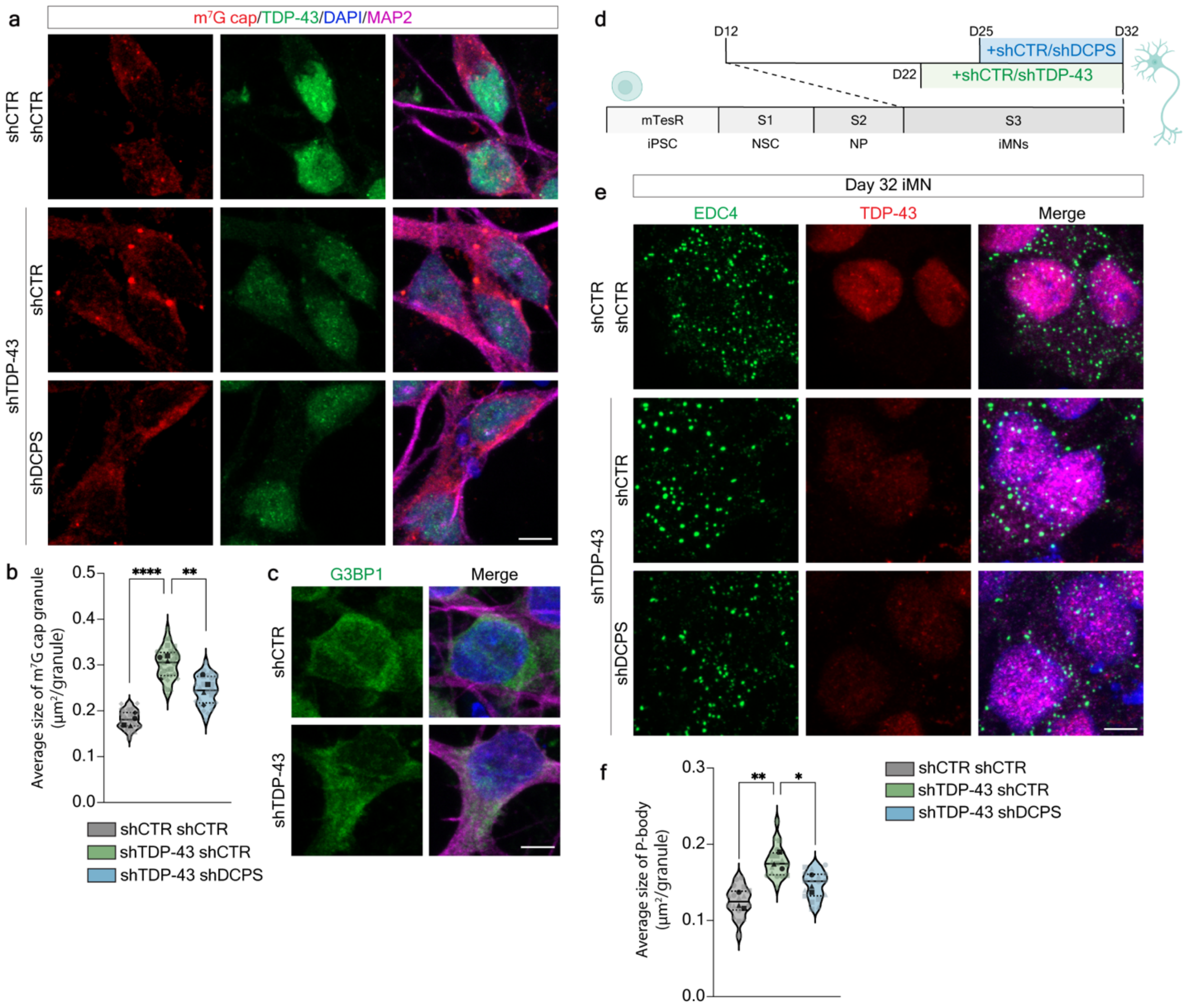


**Figure S2. DCPS reduction mitigates TDP-43 LOF-induced P-body abnormalities**

**a**, Representative immunofluorescence (IF) images of m$^7$G cap (red) and TDP-43 (green) in i$^3$Neurons. Scale bar: 5 μm. **b**, Quantification of m$^7$G-cap size in (a). The average granule size was calculated by total granule area/total granule number per imaging field (60×). Each light-colored symbol represents one imaging field. Different shapes represent biological replicates. Black symbols represent the mean value of each biological replicate. Each imaging field included at least 10 cells. The quantified imaging fields: shCTR shCTR n = 28, shTDP-43 shCTR n = 32, shTDP-43 shDCPS n = 26. Data are median (solid line) with quartiles (dashed line) from four biological replicates (black symbols). ****$P < 0.0001$, **$P = 0.0072$, by one-way ANOVA with Tukey's post hoc analysis on biological replicates. **c**, Representative IF images of G3BP1 in i$^3$Neurons with or without TDP-43 knockdown. Green, G3PB1. Magenta, MAP2. Blue, DAPI. Scale bar: 5 μm. **d**, iMN differentiation diagram with shRNA treatment timelines. **e**, Representative IF images of P-body marker EDC4 (green), TDP-43 (red), and NeuN (magenta) in iMNs differentiated from a healthy control iPSC line with different shRNA treatments. Scale bar: 5 µm. **f**, Quantification of P-body size in (e). 17 imaging fields per group were quantified. Data are median (solid line) with quartiles (dashed line) from four biological replicates. $^{**}P = 0.0059$, $^{*}P = 0.0342$, by one-way ANOVA with Tukey's post hoc analysis of biological replicates.

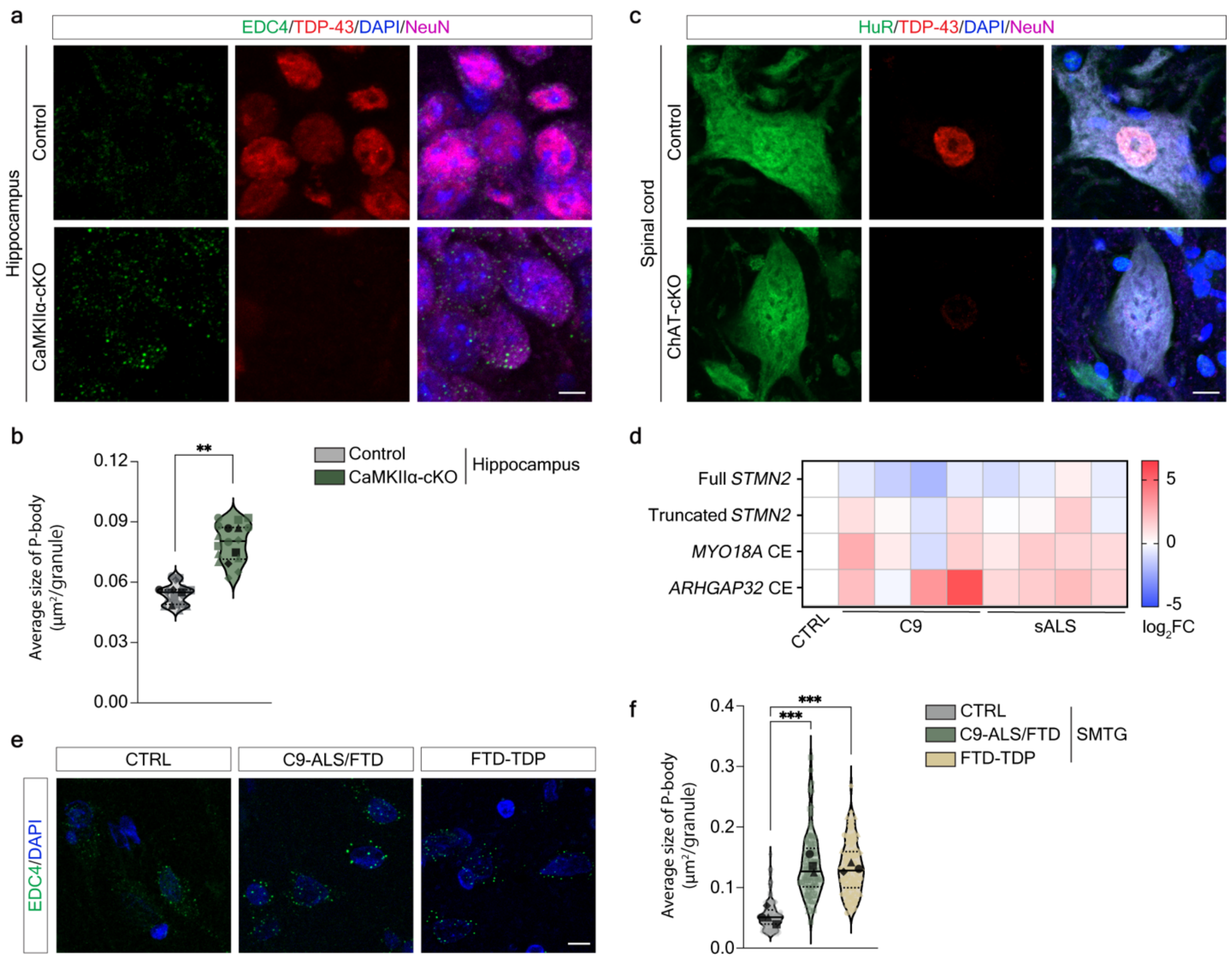


**Figure S3. P-body abnormalities in TDP-43 cKO mouse models and ALS/FTD patients**

**a**, Representative images of P-bodies in neurons from CaMKIIα$^+$ neuron-specific TDP-43 cKO mice compared to control mice at 3 months of age. Scale bar: 5 µm. **b**, Quantification of P-body size in (a). The average P-body size was calculated by total granule area/total granule number per imaging field (40×). Each light-colored symbol represents one imaging field and different shapes represent different mice. Black symbols represent the mean value of each mouse. Each imaging field included at least 70 cells. The quantified field number: Control n = 15, CaMKIIα-cKO n = 14. Data are median with quartiles from four mice. $^{**}P = 0.0018$, by two-tailed unpaired t test. **c**, Representative images of HuR staining in ChAT$^+$ motor neuron specific TDP-43 cKO mice (ChAT-cKO). Scale bar: 10 µm. **d**, Heatmap showing relative expression levels of full-length *STMN2* (Full *STMN2*) and TDP-43 LOF-induced cryptic exon (CE) isoforms (Truncated *STMN2*, *MYO18A* CE, and *ARHGAP32* CE), measured by RT-qPCR in Day-60 iMNs from *C9ORF72*-ALS/FTD (C9, n=4) and sporadic ALS (sALS, n=4) patients, compared to controls (CTRL, n=4).

The values were normalized to the control group and shown as $log_2$ fold change ($log_2FC$). **e**, Representative IF images of P-body marker EDC4 in the temporal cortex of C9-ALS/FTD, FTD-TDP patients, and age-matched non-neurological controls (CTRL). Scale bar: 10 μm. **f**, Quantification of P-body size in the temporal cortex (SMTG) of C9-ALS/FTD, FTD-TDP patients, and age-matched non-neurological controls (CTRL). The quantified cell number: CTRL n=115, C9-ALS/FTD n=93, FTD-TDP n=91. Data are median with quartiles from four cases per group. ***$P$ = 0.0001 (left), ***$P$ = 0.0002 (right), by one-way ANOVA with Tukey's post hoc analysis.

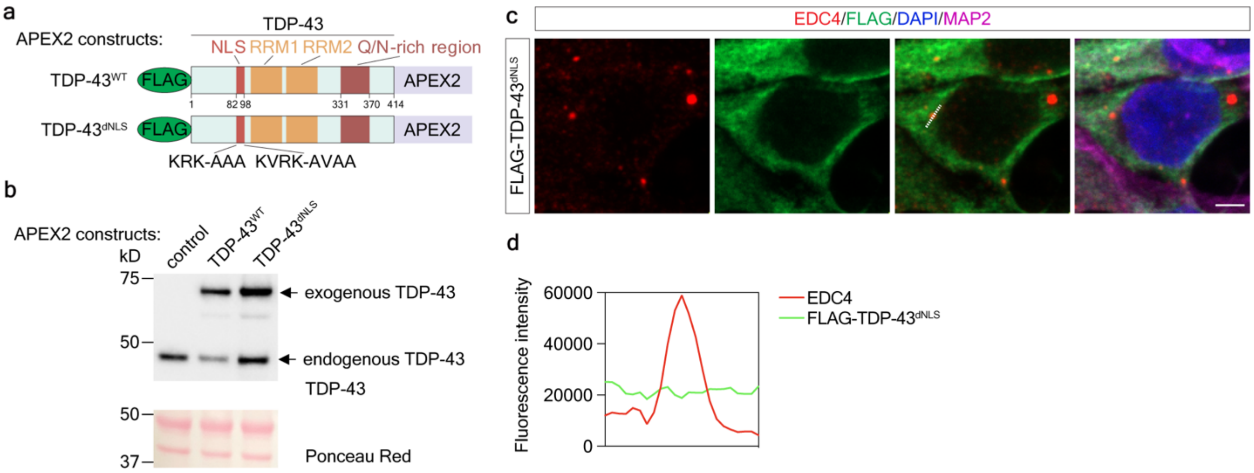


**Figure S4. APEX2 proximity labeling to identify TDP-43 interacting proteins**

**a**, Diagram of the APEX2 constructs. **b**, Western blot by anti-TDP-43 antibody to validate the expression of APEX2 constructs in $i^3$Neurons. **c**, Representative IF images of EDC4 (red) and TDP-43$^{dNLS}$ (green). Scale bar: 2.5 µm. **d**, Line profile of fluorescence intensities in (c).

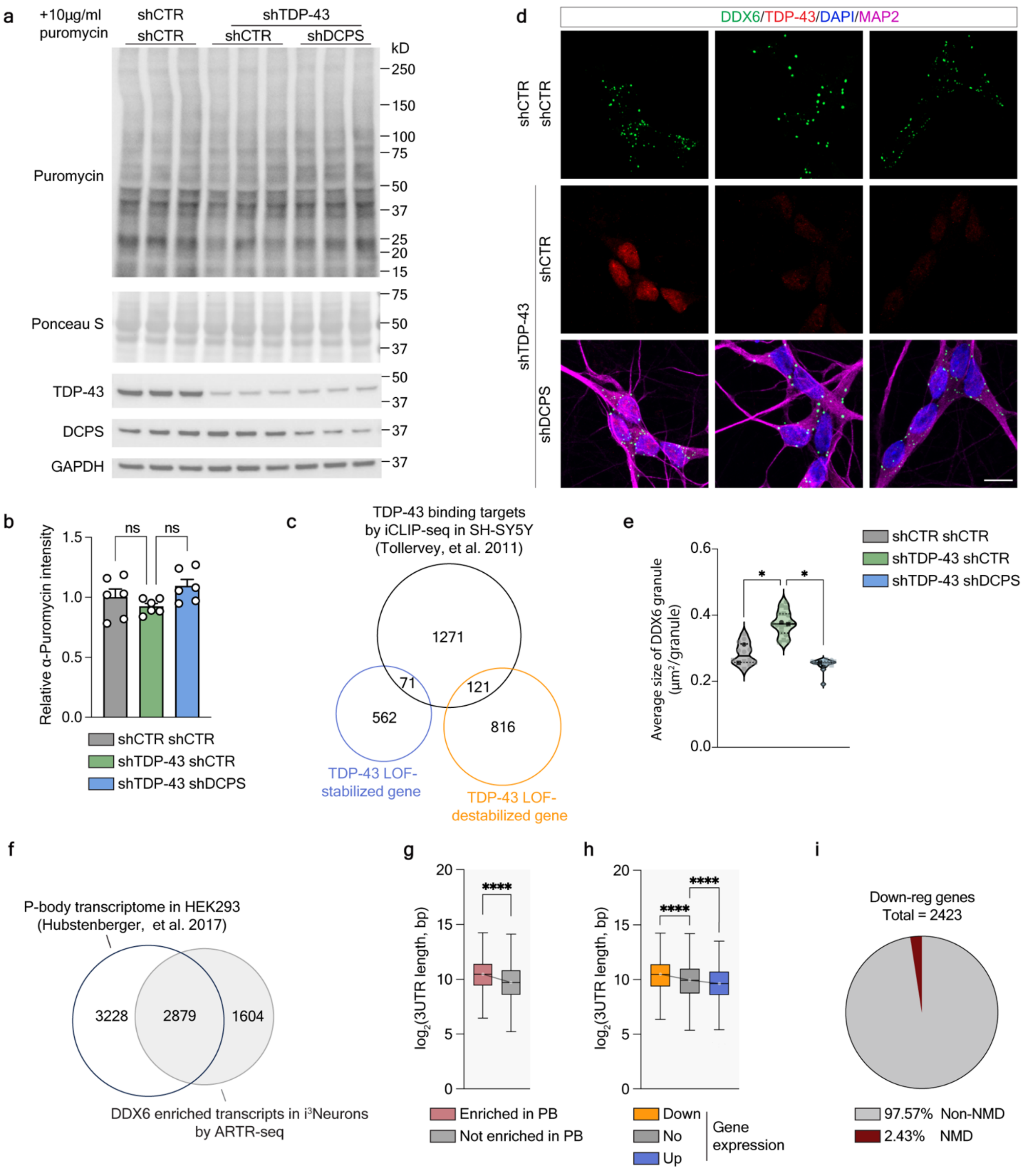


**Figure S5. TDP-43 LOF-induced P-body alteration dysregulates global RNA degradation**

**a**, Puromycin incorporation assay in i$^3$Neurons. Immunoblotting of puromycin-labeled nascent proteins using anti-puromycin antibody, with Ponceau S staining as a loading control. **b**, Quantification of the puromycin incorporation assay in (a). Data are mean ± SEM from three biological replicates with two technical replicates. $P$ = 0.5820 (ns, left), $P$ = 0.0908 (ns, right), by one-way ANOVA with Tukey's post hoc analysis. **c**, Venn diagram showing overlaps between TDP-43 binding targets by iCLIP-seq, TDP-43 LOF-stabilized and -destabilized genes. **d**, Representative images of DDX6 staining in i[3]Neurons. Scale bar: 10 µm. **e**, Quantification of DDX6-labeled P-body size in (d). Data are median with quartiles from two biological replicates. **P* = 0.0354 (left), **P* = 0.0121 (right), by one-way ANOVA with Tukey's post hoc analysis. **f**, Venn diagram showing overlap between P-body transcriptome identified by FAPS in HEK293T cells and DDX6-enriched genes revealed by ARTR-seq in i[3]Neurons. $P < 0.0001$ by Chi-square test. **g**, Box plot of 3'UTR lengths of genes enriched in P-bodies (n = 3,971) versus non-enriched genes (n = 7,709) in control neurons. Box plots indicate the interquartile range with the middle line representing the median, and the vertical lines extend to the extreme values. **** $P < 0.0001$, by Kolmogorov-Smimov test. **h**, Box plot of 3'UTR lengths of genes that were downregulated ($\log_2$FC < −0.5, adjusted $P < 0.05$, n = 2360), upregulated ($\log_2$FC > 0.5, adjusted $P < 0.05$, n = 2018), or unchanged ($|\log_2\text{FC}| \leq 0.5$ or adjusted $P \geq 0.05$, n = 6861) in neurons with TDP-43 knockdown. ****$P < 0.0001$, by Kolmogorov-Smimov test. **i**, Pie chart showing that a small proportion of genes downregulated by TDP-43 LOF are targets of nonsense-mediated decay (NMD) due to cryptic splicing.

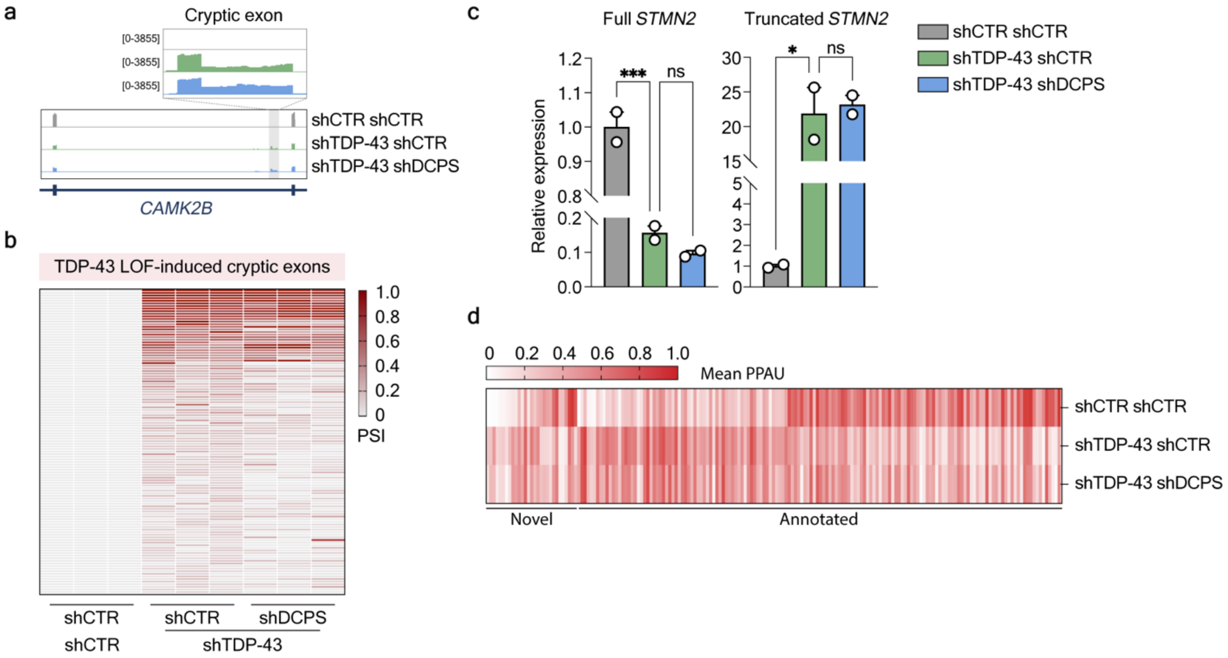


**Figure S6. DCPS reduction does not rescue cryptic splicing and APA changes induced by TDP-43 LOF**

**a**, IGV profiles of *CAMK2B* showing the cryptic exon induced by TDP-43 LOF. **b**, Heatmap shows the level of percent spliced in (PSI) of TDP-43 LOF-induced cryptic exons (n = 117) under different conditions. Cryptic exons were identified by rMATS. Data are from three biological replicates. **c**, RT-qPCR results of full-length *STMN2* isoform and cryptic-spliced isoform (truncated *STMN2*). Data are mean ± SEM from two biological replicates. ****P* = 0.0005, **P* = 0.0110, by One-way ANOVA with Tukey's post hoc analysis. **d**, Heatmap of altered alternative polyadenylation (APA) events identified by PAPA. Novel: novel APA events, n = 29. Annotated: APA events that are annotated in the reference genome, n = 154. Mean PPAU: mean polyadenylation site usage. Data are from three biological replicates.
